# Intriguing Role of Water in Plant Hormone Perception

**DOI:** 10.1101/2021.10.04.462894

**Authors:** Chuankai Zhao, Diego E. Kleiman, Diwakar Shukla

**Author notes:** Correspondence and requests for materials should be addressed to D.S.

## Abstract

Plant hormones are small molecules that regulate plant growth, development, and responses to biotic and abiotic stresses. Plant hormones are specifically recognized by the binding site of their receptors. In this work, we investigated the role of water displacement and reorganization at the binding site of plant receptors on the binding of eight classes of phytohormones (auxin, jasmonate, gibberellin, strigolactone, brassinosteroid, cytokinin, salicylic acid, and abscisic acid) using extensive molecular dynamics simulations and inhomogeneous solvation theory. Our findings demonstrated that displacement of water molecules by phytohormones contributes to free energy of binding via entropy gain and is associated with free energy barriers. Also, our results have shown that displacement of unfavorable water molecules in the binding site can be exploited in rational agrochemical design. Overall, this study uncovers the role of water molecules in plant hormone perception, which creates new avenues for agrochemical design to target plant growth and development.

## Introduction

Plant hormones are small molecules that are naturally produced *in planta* at low concentrations to regulate growth, development and stress responses.^1,2^ There are nine major classes of plant hormones that have been identified, including abscisic acid (ABA), auxin, brassinosteroid (BR), cytokinin, ethylene, giberellin (GA), jasmonic acid (JA), salicylic acid (SA) and strigolactone (SL). Generally, plants adjust their levels of plant hormones in response to a changing environment, and plant hormones act collectively to regulate a variety of responses.^1,2^ Over the last decades, exciting progress has been made in understanding different aspects of plant hormone biology, including hormone biosynthesis, transport, perception and response. In particular, the discovery of plant receptors and their crystal structures (except for ethylene) has stimulated characterization of hormone perception and signal transduction at the molecular level.^3–16^. Mechanistic understanding of hormone perception has triggered the development of synthetic agrochemicals for targeting plant receptors, which can be utilized in agricultural control of crop growth and development.^17^

Plant hormones are recognized by their receptor proteins through a host of specific protein-ligand interactions. Understanding the driving force of those protein-ligand interactions is crucial for rational design of new hormone agonists with enhanced affinity and selectivity.^20^ Water has been increasingly recognized as playing a key role in protein-ligand interactions.^21–24^ Upon ligand binding, water molecules in the cavity may be displaced, replaced, or retained to accommodate proteinligand interactions. However, due to the extremely dynamic nature of water molecules, it remains challenging to experimentally characterize structural and thermodynamic solvation properties of the binding cavity in a protein.^22,23,25^ Explicit solvent molecular dynamics (MD) simulations capture the motion of proteins in a solution environment, making it an ideal method for addressing the challenges in studying water in the buried regions of protein.^26^ MD simulations have also been used extensively in studying detailed protein-ligand binding mechanisms.^27–30^ In recent years, a range of computational tools based on inhomogeneous solvation theory (IST) have emerged as powerful techniques to investigate solvation structural and thermodynamic properties of binding cavity. Based on these tools, solvation thermodynamic information can be exploited in predicting ligand binding affinity and rational ligand optimization.^31–33^

In this work, we performed large-scale MD simulations (aggregate ~786 μs) to investigate the role of water on the perception of eight major classes of plant hormones (auxin, JA, GA, SL, BR, cytokinin, SA, and ABA) by their receptors in *Arabidopsis thaliana* (Figure 1).^30,34–37^ Markov state model (MSM) analysis was used to characterize the thermodynamics associated with the binding processes.^38,39^ By analyzing the binding pathways, we assessed the key role of water in plant hormone binding. Furthermore, we characterized the solvation structural and thermodynamic properties of the eight plant receptors by analyzing MD trajectories on *apo* proteins via IST-based hydration site analysis (Supplementary Table S1). Based on these data, we analyzed the properties of water molecules upon the binding of plant hormones, and quantified their contribution to overall free energy of binding. Finally, we characterized the thermodynamic properties of unfavorable water molecules that are retained in the bound complexes. We showed that excluding such unfavorable water molecules by optimizing a ligand can lead to enhanced ligand binding affinity. Overall, our findings elucidate the role of water in perception of plant hormones, and create potential avenues for agrochemical discovery to target plant receptors.

**Figure 1:**
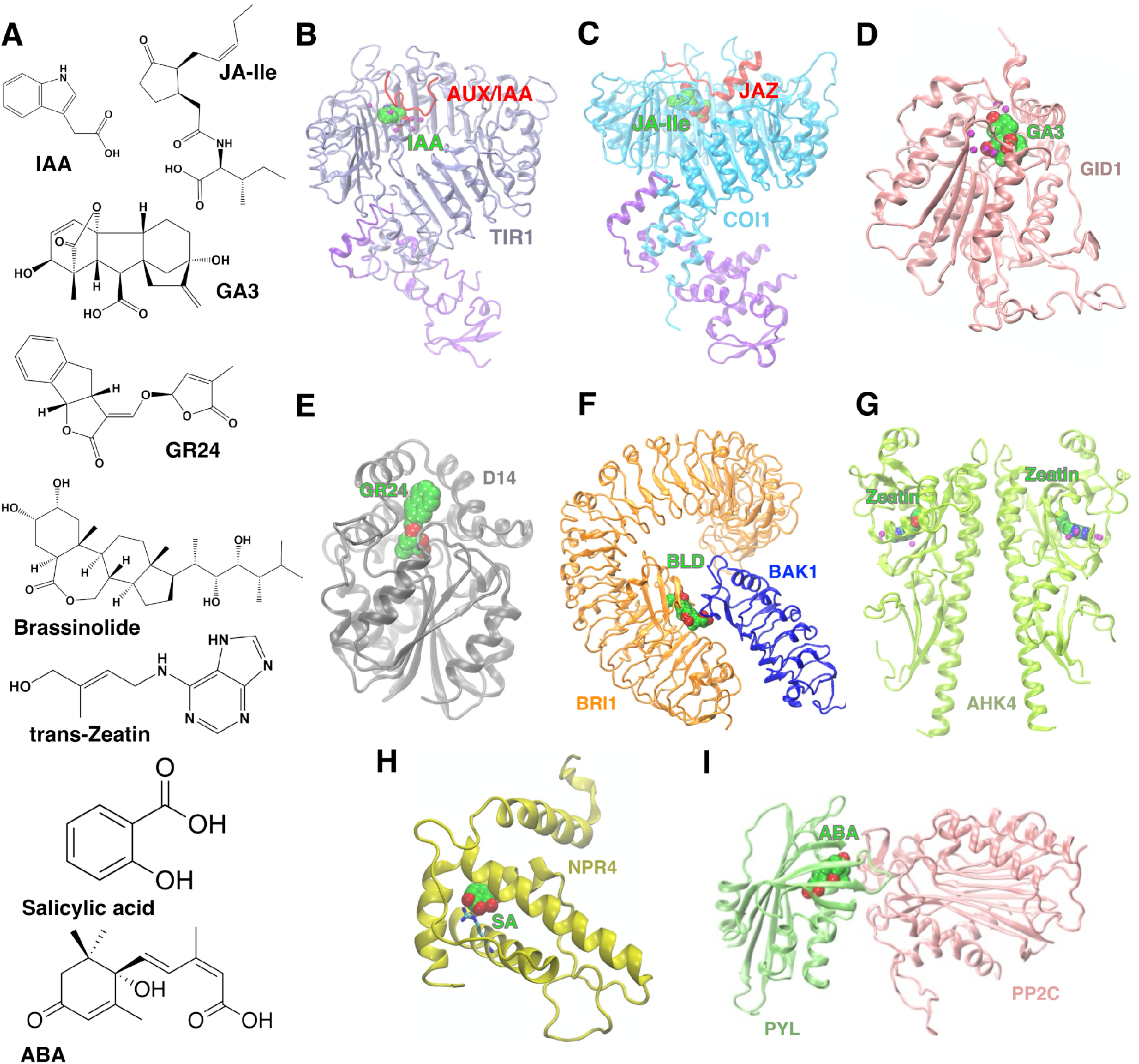
Plant hormones and their receptors. (A) Molecular structures of eight major classes of plant hormones. (B) TIR1-AUX/IAA for auxin (PDB ID: 2P1Q^3^), (C) COI1-JAZ for jasmonate (PDB ID: 3OGL^5^), (D) GID1 for gibberellin (PDB ID: 2ZSH^4^), (E) D14 for strigolactone (PDB ID: 5DJ5^11^), (F) Extracelluar domains of BRI1 and BAK1 for brassinosteroid (PDB ID: 4LSX^18^), (G) extracelluar sensor domain of AHK4 for cytokinin (PDB ID: 3T4L^10^), (H) NPR4 and salicylic acid (PDB ID: 6WPG^13^), (I) PYL2-PP2C and ABA (PDB ID: 4LA7^19^).

## Results

### Long timescale MD simulations capture the pathway of plant hormone binding and the relevance of water in this process

First, we sought to characterize how water in the binding sites affect phytohormone binding processes. We performed large timescale MD simulations (aggregate ~786 μs) to capture the binding of IAA, JA-Ile, GA3, trans-zeatin, and SA to their respective receptors, followed by MSM analysis on the binding trajectories (Supplementary Methods, Supplementary Figure S1, Supplementary Tables S2-7). Using MSMs, the pathway for the binding of plant hormones and the associated thermodynamics and kinetics can be quantified.^38,39^ We also obtained the data for the binding of BLD, GR24 and ABA from past studies (BLD from ref. 30, GR24 from ref. 34, and ABA from ref. 35). We reported the free energy landscapes that describe the overall binding of plant hormones as well as the pathways for these processes identified from transition path theory (Supplementary Figures S2-9).

Employing the resulting MSMs, we interrogated the role of waters in the binding processes. The solvation shell of a phytohormone accounts for the interfacial waters that bridge the pocket-ligand interactions during hormone-receptor complexation. These solvent molecules must be either displaced or reorganized to accommodate the ligand-protein interactions. These processes can pose significant energy barriers for ligand binding. For this reason, we assessed the role of water molecules in hormone binding by computing the free energy landscape of ligand hydration against the binding coordinate resolved by tICA (Fig. 2). These plots show that the binding process is correlated with ligand desolvation. The presence of a free energy barrier along the solvation coordinate shows that water displacement is a key process for hormone-receptor complexation. Due to the diversity of the ligand-receptor systems analyzed, stark contrasts are also evidenced in these free energy landscapes. For example, the plot for SA-NPR4 shows a funnel-like landscape with a large energy barrier (~8 kcal/mol) between states 1 and 2. As the binding process of SA continues, the ligand loses all the waters in its hydration shell, mostly due to the formation of hydrophobic interactions with pocket residues. On the other hand, systems such as GA3-GID1, which present several bridging waters, show lower energy barriers (~2 kcal/mol) along the binding pathway. It is also observed that some of the systems (e.g., trans-Zeatin-AHK4 and ABA-PYL2) show free energy minima that are not associated with the main binding pathway and indicate alternative ligand desolvation paths. Such minima are generally associated with off-target binding.

**Figure 2:**
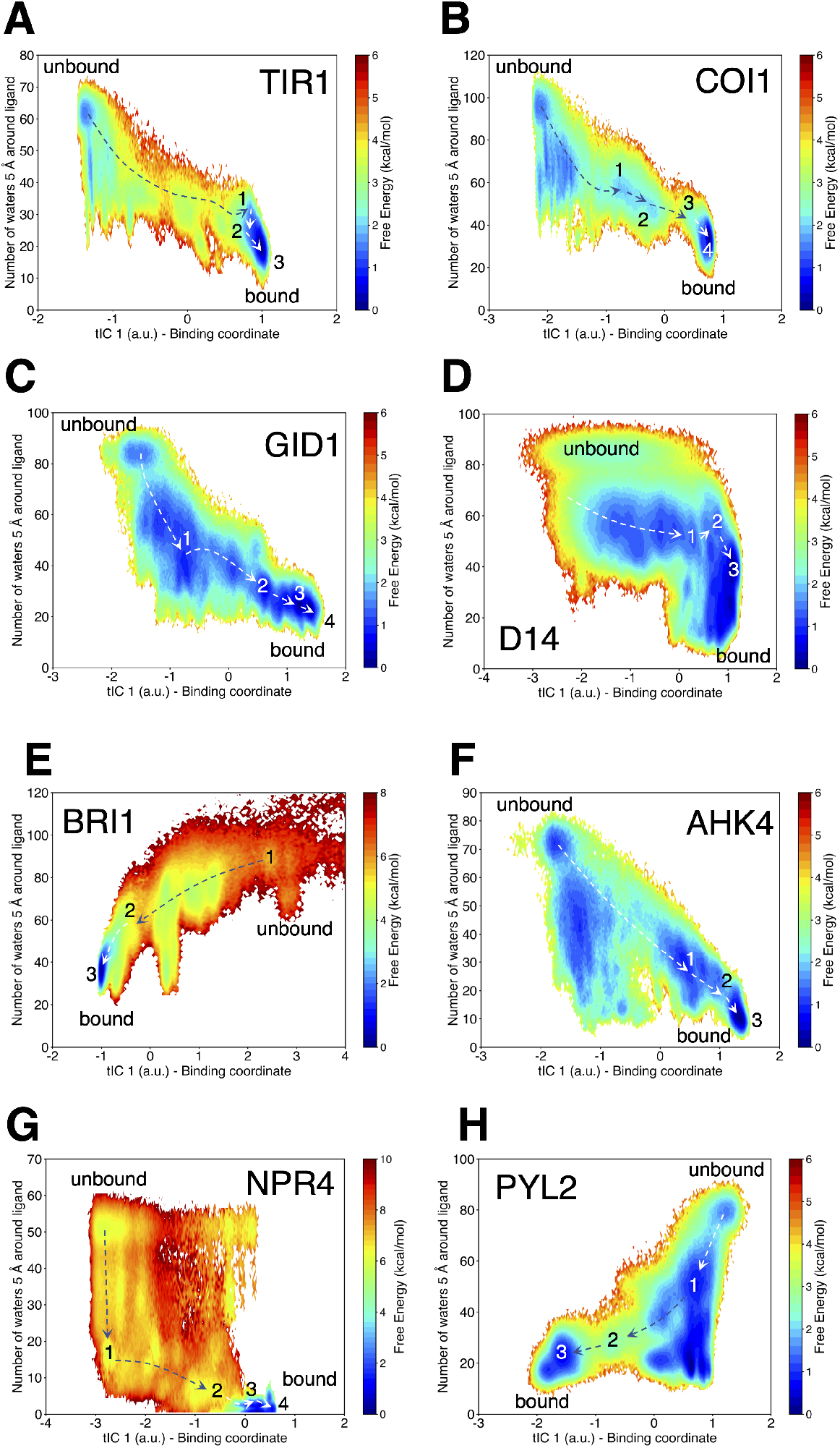
Free energy landscapes for ligand solvation of plant hormones upon receptor binding. (A) IAA binding to TIR1, (B) JA-Ile binding to COI1, (C) GA3 binding to GID1, (D) GR24 binding to D14, (E) BLD binding to BRI1, (F) trans-zeatin binding to AHK4, (G) SA binding to NPR4, and (H) ABA binding to PYL2. The *x* coordinate represents the slowest degree of freedom from time-lagged independent component analysis, which captures the binding process. The *y* coordinate is the number of water molecules within 5 Å of ligand (approximately first and second solvation shells).

Although free energy landscapes allow us to assess the relevance of the solvent in the binding process in terms of the water displacement barrier, several important factors remain unaccounted for by this methodology. Namely, the distribution of hydration clusters in the binding site and their thermodynamic properties need to be evaluated individually to determine if their displacement would further stabilize the bound complex. This is the focus of the following sections.

### Solvation structural and thermodynamic properties of *apo* plant receptors

In this section, we focused on characterizing the structural and thermodynamic properties of hydration sites in the hormone binding pockets to resolve the detailed energetic contributions of waters in ligand-receptor complexation. In order to investigate the solvation pattern of the binding site of *apo* plant receptors, we performed 100 ns explicit solvent MD simulations on the eight receptors. The crystal structures of the plant receptors in the hormone-bound conformations were used and conformational restraints on protein backbone were applied in MD simulations (Supplementary Methods). IST-based hydration site analysis (HSA) was then used to analyze the trajectories from MD simulations in order to characterize solvation structural and thermodynamic properties of the binding cavities.^31–33^ Hydration sites define the centers in the binding pocket where water resides with the highest probabilities. Using IST, the enthalpic (including both protein-water and water-water interaction energy, denoted as *E_sw_* and *E_ww_*) and entropic properties (excess entropy relative to bulk water — *TS^e^)* of water molecules in hydration sites can be estimated. Furthermore, structural properties of water molecules such as the number of hydrogen bonds with protein 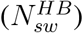 and neighboring water molecules 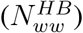 can be determined. We identified the hydration sites in the eight plant receptors, which are numbered according to water occupancy probabilities (*f_o_*) (Figure 3, Supplementary Tables S8-15). Based on the local environment, the hydration sites were classified into apolar (A), polar (P), and charged (C) sites. In addition, the sites were classified into either favorable (F) or unfavorable (U) by comparing *E_tot_* with bulk water. Lastly, these sites were also classified into either enhanced (En, favorable water-water interactions) and frustrated (Fr, unfavorable water-water interactions) by comparing *E_ww_* of each site and bulk water (Supplementary Methods, Supplementary Figure S10).

**Figure 3:**
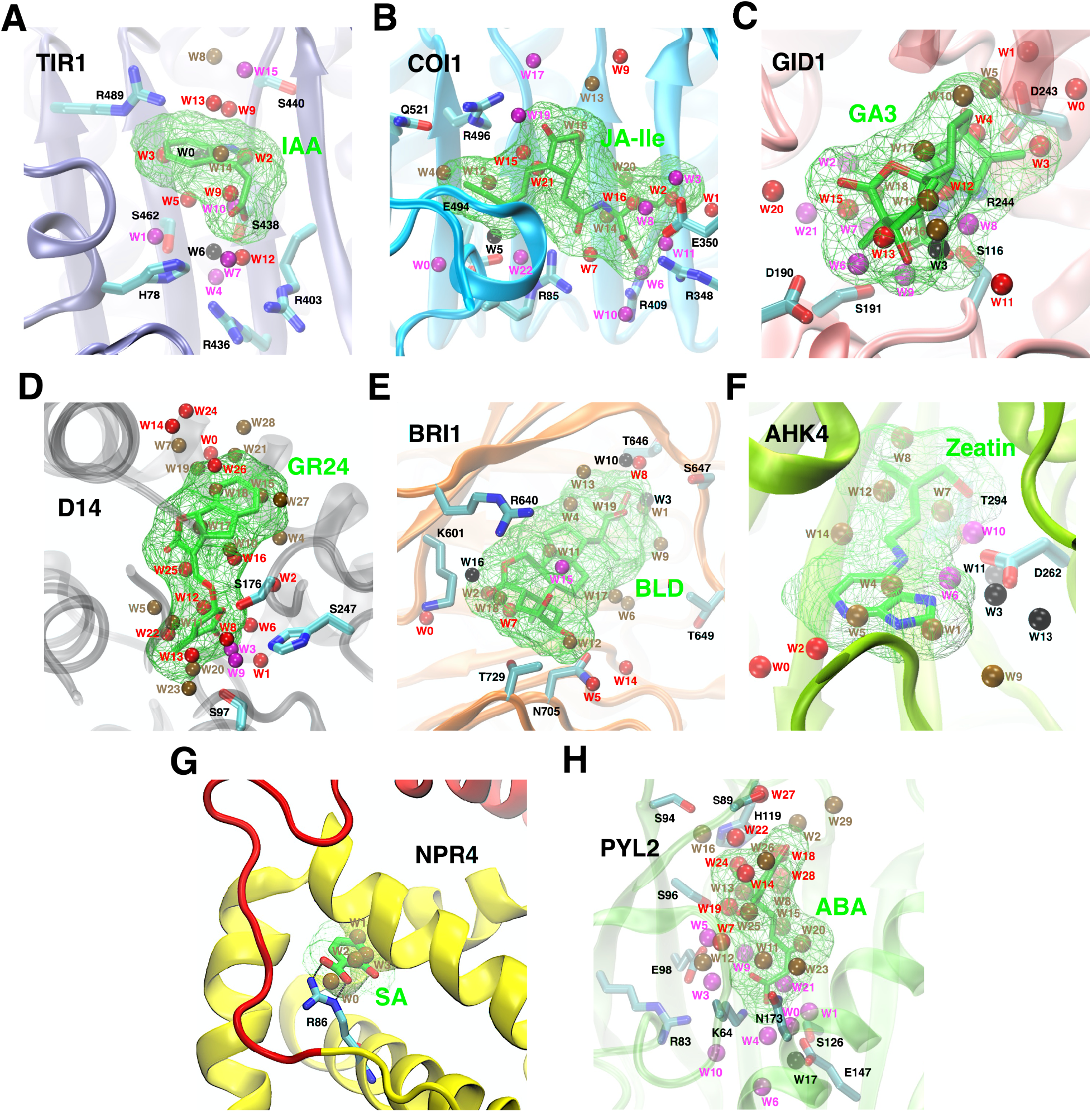
Visualization of hydration sites at the binding sites of (A) TIR1 in iceblue, (B) COI1 in cyan, (C) GID1 in pink, (D) D14 in grey, (E) BRI1 in orange, (F) AHK4 in lime, (G) NPR4 in yellow/red, and (H) PYL2 in green. The plant hormones in the bound poses and their envelopes (probe radius of 1.5 Å) are shown in green. The polar and charged residues at the binding sites are shown in cyan. Hydration sites are numbered based on their occupancy and colored according to their energetic states: En.F (red), En.U (ochre), Fr.F (magenta) and Fr.U (black).

In total, we have identified 159 hydration sites in the eight receptors (A: 22.64%, P: 44.03%, C: 33.33%, Table 1). For five of the receptors, we show that our predicted locations of the hydration sites match (up to 2 Å) with most of the water locations reported in the hormone-bound crystal structures (Supplementary Figure S11). For all the eight receptors, there are generally more enhanced sites than frustrated sites (Figure 4). For TIR1, COI1, GID1, and PYL2, due to the presence of several charged residues at the binding site, there are enhanced populations of frustrated sites and frustrated sites are generally favorable (Figure 4A, B, C, H). In contrast, the hydration sites in D14, BRI1, AHK4, and NPR4 are dominantly enhanced due to overall hydrophobic environment in the binding cavity (Figure 3D, E, F, G and Figure 4D, E, F, G). In addition, the number of En.U sites is greater than any other type of hydration site for D14, BRI1, AHK4, NPR4, and PYL2 (Figure 4D, E, F, G, H). Overall, MD simulations and hydration site analysis have provided detailed characterization of solvation properties at the binding site of plant receptors.

**Figure 4:**
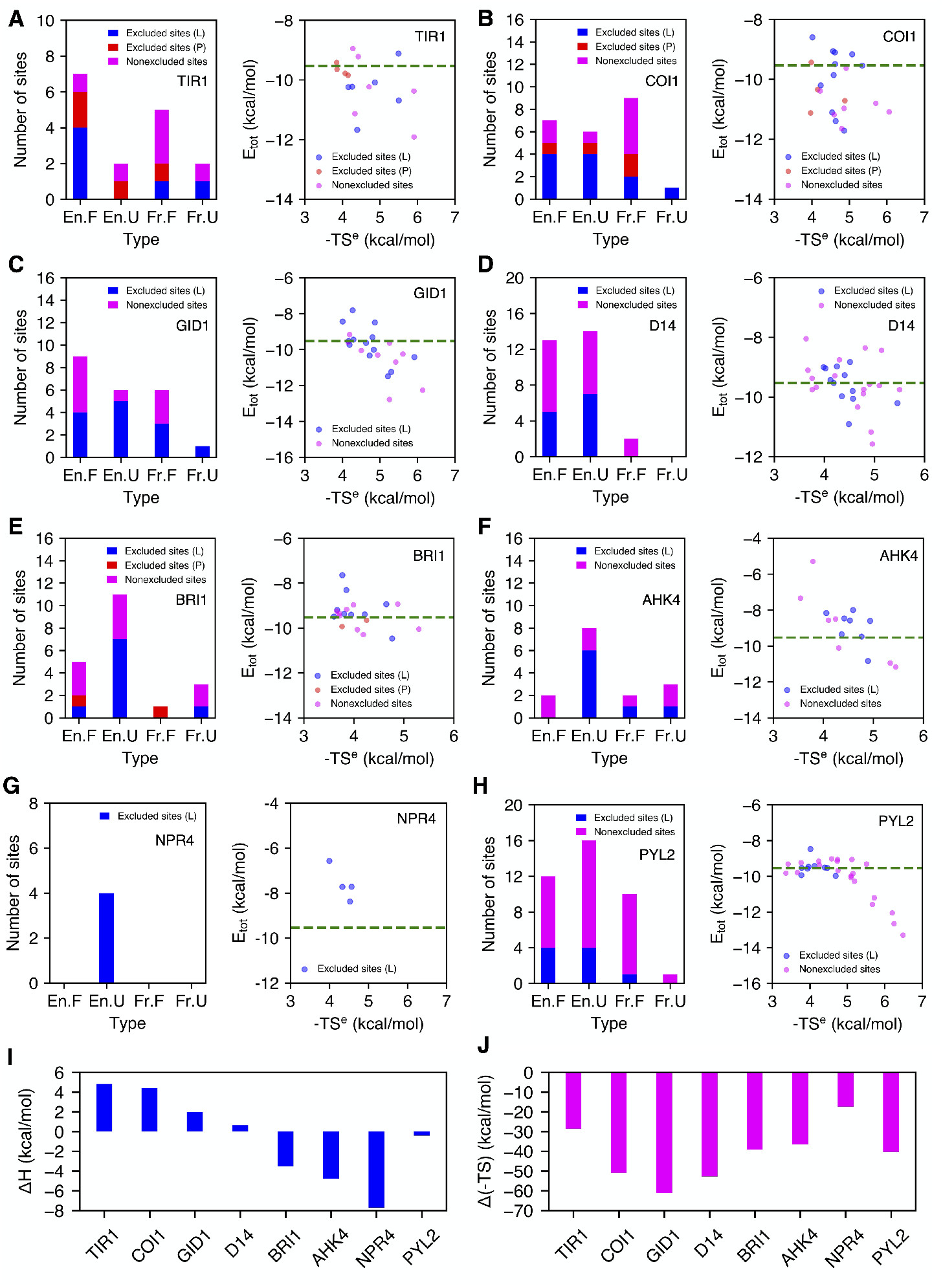
Displacement of water from hydration sites at the binding cavities. (Left) number of hydration sites (En.F, En.U, Fr.F and Fr.U) that are displaced upon the binding of plant hormones and co-receptors, and (right) *E_tot_* and -TS^*e*^ of the hydration sites for (A) TIR1, (B) COI1, (C) GID1, (D) D14, (E) BRI1, (F) AHK4, (G) NPR4, and (H) PYL2. The hydration sites displaced by plant hormones (L) and co-receptors (P) are shown in blue and red, and the retained sites are shown in magenta. Overall (I) enthalpic and (J) entropic contribution of water displacement to free energy of binding of plant hormones to the eight receptors.

**Table 1:** The average thermodynamic and structural quantities for all the hydration sites identified from the binding cavity of plant hormone receptors (energy unit: kcal/mol). Data for pure water is included for comparison.

| type | count | $f_o$ | $E_{sw}$ | $E_{ww}$ | $E_{tot}$ | $E_{ww}^{nbr}$ | $N_{nbr}$ | $f_{enc}$ | $N_{sw}^{HB}$ | $N_{ww}^{HB}$ | $f_{ww}^{HB}$ | $N_{ww,lost}^{HB}$ | frac. |
| --- | --- | --- | --- | --- | --- | --- | --- | --- | --- | --- | --- | --- | --- |
| pure water | - | - | 0.00 | -9.53 | -9.53 | -1.36 | 5.26 | 0.00 | 0.00 | 3.33 | 0.63 | 0.00 | - |
| A.En.F | 11 | 0.40 | -2.15 | -7.92 | -10.1 | -1.63 | 4.21 | 0.20 | 0.01 | 3.17 | 0.75 | 0.16 | 0.64 |
| A.En.U | 24 | 0.39 | -1.97 | -7.01 | -8.98 | -1.72 | 3.46 | 0.34 | 0.01 | 2.62 | 0.76 | 0.71 | 0.67 |
| A.Fr.U | 1 | 0.30 | 0.16 | -5.45 | -5.29 | -1.10 | 4.00 | 0.24 | 0.07 | 2.35 | 0.59 | 0.98 | 0.00 |
| C.En.F | 11 | 0.61 | -6.06 | -4.56 | -10.6 | -1.64 | 2.64 | 0.50 | 1.20 | 2.00 | 0.76 | 1.33 | 0.36 |
| C.En.U | 11 | 0.65 | -4.01 | -4.67 | -8.68 | -1.86 | 2.23 | 0.57 | 0.89 | 1.69 | 0.75 | 1.66 | 0.73 |
| C.Fr.F | 26 | 0.64 | -9.19 | -1.78 | -11.0 | -0.83 | 2.65 | 0.50 | 1.47 | 1.55 | 0.58 | 1.78 | 0.20 |
| C.Fr.U | 5 | 0.53 | -7.23 | -1.89 | -9.12 | -0.96 | 2.60 | 0.51 | 1.01 | 1.54 | 0.64 | 1.79 | 0.40 |
| P.En.F | 29 | 0.52 | -4.64 | -5.49 | -10.1 | -1.69 | 2.83 | 0.46 | 0.95 | 2.14 | 0.76 | 1.19 | 0.38 |
| P.En.U | 28 | 0.42 | -3.20 | -5.58 | -8.78 | -1.75 | 2.70 | 0.49 | 0.64 | 1.99 | 0.75 | 1.36 | 0.50 |
| P.Fr.F | 8 | 0.47 | -6.25 | -4.22 | -10.5 | -1.15 | 3.14 | 0.40 | 1.10 | 2.00 | 0.64 | 1.33 | 0.38 |
| P.Fr.U | 5 | 0.48 | -5.06 | -4.13 | -9.19 | -1.22 | 2.82 | 0.46 | 1.15 | 1.72 | 0.60 | 1.61 | 0.60 |
| Undisplaced | 86 | 0.53 | -5.52 | -4.40 | -9.93 | -1.37 | 2.88 | 0.46 | 0.97 | 1.97 | 0.69 | 1.36 | - |
| Displaced | 73 | 0.48 | -3.86 | -5.60 | -9.46 | -1.65 | 2.99 | 0.43 | 0.60 | 2.21 | 0.75 | 1.13 | - |

### Displaced water molecules from enhanced hydration sites contribute to binding affinity via entropy gain

We further sought to understand how water molecules in the pocket affect the binding of plant hormones. In this work, we have focused our analysis on one member of endogenous or synthetic analogs of natural plant hormones (IAA for auxin, JA-Ile for JA, GA3 for GA, synthetic analog of SL (GR24), brassinolide (BLD) for BR, trans-zeatin for cytokinins, SA, and ABA; Figure 1A). We analyzed the displacement of water molecules by the binding of these ligands, by assuming that any water in the hydration sites that are within 1.5 A of their bound poses would be excluded. In this way, we identified all the hydration sites to be displaced upon the binding of plant hormones, the functional groups in the ligands responsible for displacing certain types of hydration sites (Figure 3), and their contributions to the free energy of binding (Figure 4). To show the exclusion of water molecules along the ligand binding processes, we showed *apo* hydration sites over the binding intermediate states captured from our MD simulations and MSM analysis (Supplementary Figures S2-9). Generally, exclusion of favorable hydration sites would lead to a larger free energy barrier to ligand binding.

Overall, 73 out of 159 hydration sites in all the eight receptors are displaced upon the binding of plant hormones. 60 displaced sites belong to En categories (82.20%) and 43 sites belong to U categories (58.90%). Generally, water molecules in enhanced and unfavorable hydration sites are more likely to be displaced upon the binding of plant hormones. The fractions of displaced hydration sites in apolar, polar and charged sites are 64.22%, 44.29%, and 36.21% respectively. The displaced hydration sites tend to have relatively weaker protein-water interaction compared to the undisplaced sites (Table 1).

The enthalpic contribution of displaced water molecules to overall free energy of binding depends on thermodynamic properties of water molecules. Other than TIR1, the majority of water molecules displaced by the ligands are from En.U hydration sites. Displacing En.U water molecules leads to a favorable contribution to enthalpy change in binding, in addition to their favorable entropic contribution. En.U hydration sites are generally displaced by a hydrophobic alkyl chain or a bulky ring of the ligands (Figure 3B-H). Strikingly, 7 out of 9 displaced hydration sites in BRI1, 6 out of 8 displaced hydration sites in AHK4, and all 4 displaced sites in NPR4 are En.U (Figure 4E, F, G). The displacement of one Fr.U hydration site in BRI1 and AHK4 further leads to negative binding enthalpy changes. Overall, expelling water molecules from the binding site of BRI1 and AHK4 contribute to free energy of binding both enthalpically and entropically (Figure 4I, J).

The second major type of displaced hydration site is En.F site. There are 4-5 water molecules from En.F sites being expelled into the bulk for TIR1, COI1, GID1, D14, and PYL2. Displacement of such favorable sites result in enthalpic penalty for ligand binding, which requires to be compensated by favorable protein-ligand interactions. Generally, En.F sites are displaced by a bulky ring or a polar functional group in the ligands (Figure 3A-E, H).

Fr.F is the third type of hydration site being displaced as seen in TIR1, COI1, GID1, AHK4, and PYL2. Fr.F is usually observed around the charged residues in the binding cavity (Figure 3A-F, H). Displacing Fr.F site can result in significant enthalpic penalty to ligand binding, which requires to be compensated by strong protein-ligand interactions. A common observation is that Fr.F sites are displaced by the carboxylate group of the ligands (Figure 3A-C, H) which form electrostatic interactions with charged or polar residues. Taken altogether, displaced water molecules are mostly from En.U hydration sites, and contribute to free energy of binding via gain of entropy.

### Identification of unfavorable water molecules in the bound complex for agrochemical optimization

After the binding of plant hormones, water molecules from some *apo* hydration sites, including unfavorable hydration sites (En.U or Fr.U), may remain in the bound complex (Figure 4). These water molecules can either be stabilized or destabilized by the presence of plant hormones in the binding site. If a water molecule with unfavorable *E_tot_* exists in the bound complex, optimizing the ligand to exclude such water upon binding can lead to further enthalpy gain for ligand binding. We further sought to determine the thermodynamic properties of water molecules in the bound complex. For each receptor, we extracted the ligand-bound conformation with water molecules present in the binding site from our binding MD simulations. Then, we performed 100 ns MD simulations starting from these snapshots followed by hydration site analysis to determine the hydration sites in the bound complex (denoted as *holo).* We note that the *holo* hydration sites may not reproduce all the nonexcluded *apo* hydration sites, since the pocket environment in the chosen bound configuration may deviate from the receptor conformation in *apo* simulations. We therefore focused on characterizing the changes in enthalpy and entropy for the conserved sites captured in *apo* and *holo* simulations, particularly for *apo* unfavorable hydration sites (Figure 5A,B and Supplementary Figure S12-S17). While some of the unfavorable *apo* hydration sites are indeed stabilized upon the binding of plant hormones, we identified a few buried unfavorable hydration sites in COI1 (site 14 and 20, Figure 5A,B), D14 (site 5, Supplementary Figure S14), and PYL2 (sites 2 and 29, Supplementary Figure S17) that are further destabilized by the bound ligands.

**Figure 5:**
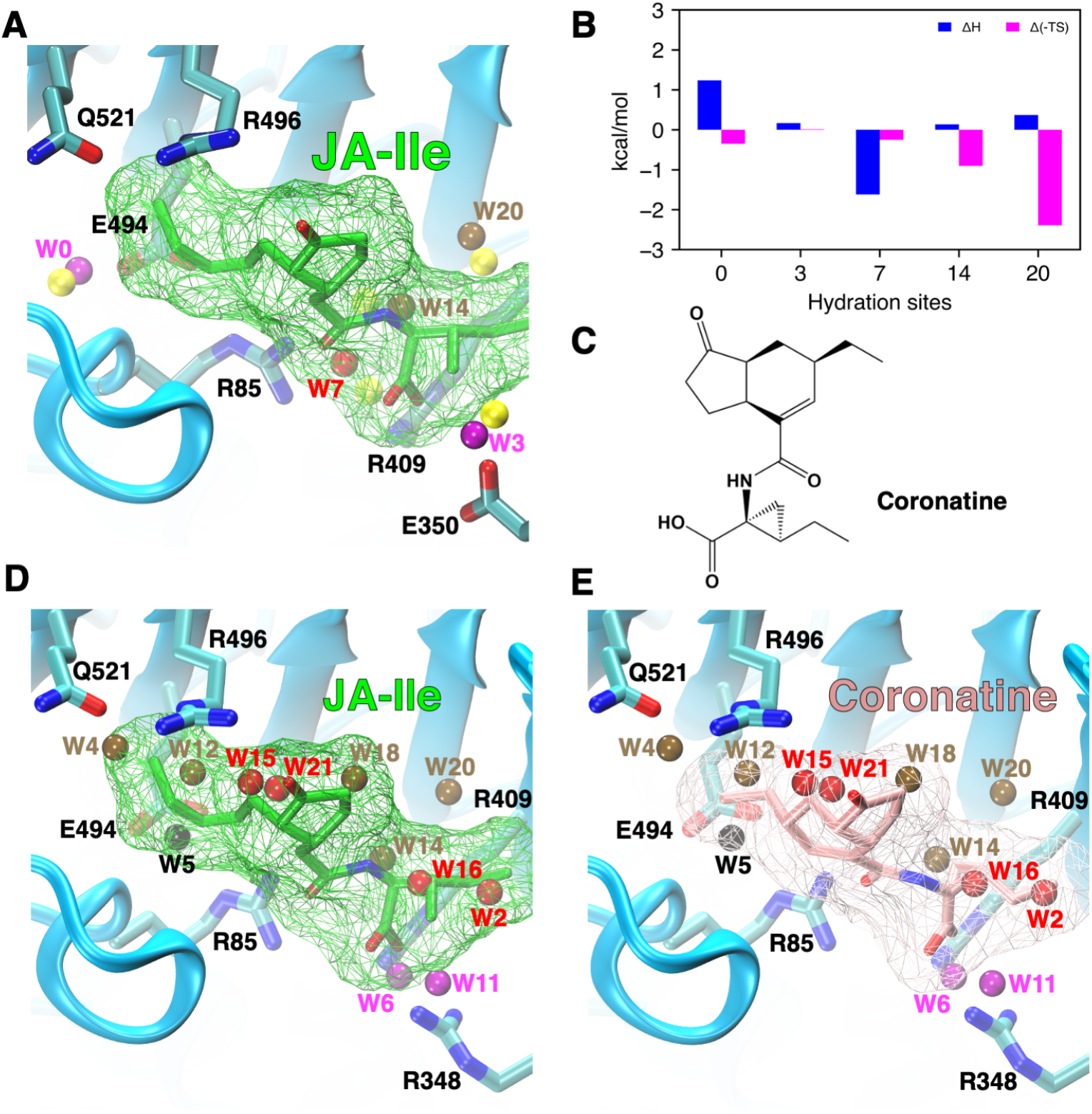
Molecular origin of enhanced affinity of JA analog coronatine. (A) Overlay of the conserved apo and holo hydration sites in the binding site of COI1 upon the binding of JA-Ile, and (B) the changes in enthalpy and entropy of conserved hydration sites. (C) Molecular structure of coronatine. (D-E) Comparison of the exclusion of water molecules upon the binding of (D) JA-Ile and (E) coronatine to COI1.

The presence of the unfavorable hydration site 14 in the bound complex of COI1 and JA-Ile can partially explain the 10-fold enhanced affinity of JA analog coronatine relative to JA-Ile.^5^ Based on the crystal structure of the coronatine-bound complex, the cyclopropane ring present in coronatine would completely exclude the unfavorable hydration site 14, thereby contributing to enthalpy gain in binding affinity. In addition, the cyclohexene ring results in enhanced hydrophobic interaction between COI1 and the ligand to compensate for the enthalpy loss due to the displacement of favorable hydration sites 15 and 21 (Figure 5E). However, the shorter alkyl chain in coronatine leaves the unfavorable hydration site 4 in the bound complex, which is displaced by JA-Ile. For both JA-Ile and coronatine, unfavorable site 20 remains in the bound complex, which can potentially be excluded by further ligand optimization to enhance binding affinity.

Moreover, HSA could partially explain why the PYL2 antagonist pyrabactin (Figure S18) binds PYL2 with a slightly lower affinity compared to ABA.^40^ Based on the crystal structure of the PYL2-pyrabactin complex (PDB ID: 3NJ0^41^) the sulfonamide group in pyrabactin would displace hydration site 9 (Fr.F), unlike ABA (Figure S18). This represents an enthalpic penalty which is not compensated by a protein-ligand interaction. Additionally, pyrabactin would not displace waters in sites 13 (En.U), 18, and 24 (both En.F). The retention of favorable sites provides an enthalpic benefit to pyrabactin binding, while the retention of site 13 represents an entropic loss. On the other hand, the pyridyl group in pyrabactin displaces hydration site 15 (En.U) and establishes hydrophobic interactions with nearby residues, which enhances its binding affinity. Also, pyrabactin would not displace site 21 (Fr.F), but in the case of ABA, the carboxylate group displaces this water to form a strong interaction with LYS64. Neither ABA nor pyrabactin displace sites 2 or 29, which could potentially be excluded through ligand optimization and lead to further complex stabilization.

Overall, these results have shown that displacement of unfavorable water molecules in the binding site of plant receptors can further enhance ligand binding affinity. Moreover, the results also suggest a pathway to agrochemical design by optimizing ligands to exclude certain hydration sites.

## Discussion

In this study, we present an in-depth computational investigation of the role of water on the binding of eight major classes of plant hormones to their receptors. The hydrophobic pocket results in overall enhanced local water structure in the cavity, and water molecules may be either favorable or unfavorable relative to bulk which depends on protein-water and water-water interactions. The presence of charged residues in the pocket mostly leads to frustrated and favorable local water structure. We have shown that the binding of plant hormones mostly displace enhanced hydration sites into the bulk, and displaced hydration sites tend to have weaker protein-water interactions. We have also identified the functional groups that are responsible for displacing different types of hydration sites. En.U sites are generally displaced by hydrophobic groups in the hormones, and En.F sites are displaced by bulky hydrophobic groups such as aromatic rings. On the other hand, Fr.F sites are preferably excluded by charged or polar functional groups in the ligand which can form favorable interaction with protein to compensate for the enthalpy loss of favorable water molecules.

Our results have demonstrated that displacement and reorganization of water molecules upon the binding of plant hormones have a significant contribution to overall free energy of binding. For BRI1, AHK4, and NPR4, the displacement of unfavorable water molecules lead to enthalpy gain of hormone binding. For the rest of the plant receptors, the displacement of water molecules results in some enthalpy loss and therefore unfavorable enthalpic contribution to free energy of binding, due to the exclusion of favorable water molecules. The water molecules in the binding cavity have significant entropy loss compared to bulk water. Overall, the displacement of water molecules contribute to free energy of binding via gain of entropy of water molecules.

Solvation structural and thermodynamic properties of the binding cavity of plant receptors can be potentially exploited in future agrochemical discovery and optimization. We have shown that water molecules from unfavorable hydration sites can remain in the ligand-bound complex, and some of them can be further destabilized by the binding of plant hormones. Enhanced binding affinity can be achieved by redesigning a ligand to displace such unfavorable water molecules. We have utilized the solvation thermodynamic data to explain the enhanced affinity of JA analog coronatine relative to JA-Ile. We have also used these results to rationalize the lower affinity of pyrabactin to PYL2 when compared to ABA. Recent studies have also demonstrated that manipulation of water network in the receptor binding site during lead optimization can result in significant increase of ligand binding affinity.^21–23^ Also, methods are being developed to incorporate solvation thermodynamic information into binding affinity evaluation in a virtual screening process.^42^

In conclusion, our results provide new insights into the fundamental mechanism of plant hormone perception, and suggest that water plays an important role in the binding of plant hormones. We expect that our findings can help advance our understanding of the role of water in protein-ligand binding, and create new avenues for designing hormone agonists with enhanced affinity.

## Supporting information

Supplementary Methods, Images, Tables and Results

## Acknowledgements

The authors acknowledge the support from the Blue Waters sustained-petascale computing project, which is funded by the National Science Foundation (awards OCI-0725070 and ACI-1238993) and the state of Illinois. D.S. acknowledges the support from Foundation for Food and Agriculture Research via the New Innovator Award in Food & Agriculture Research. C.Z. acknowledges the support by 3M Corporate Fellowship and Glenn E. and Barbara R. Ullyot Graduate Fellowship from the University of Illinois at Urbaba-Champaign. C.Z. and D.E.K. would like to thank Dr. Zahra Shamsi for setting up MD simulations of the binding of trans-zeatin to AHK4 and Ph.D. candidate Jiming Chen for providing data on the D14-GR24 system.

## Author Contributions

C.Z. and D.S. designed the study. C.Z. and D.E.K. performed the MD simulations, analyzed the data and wrote the manuscript with input from D.S.

## Addtional Information

**Supplementary Information** accompanies this paper online.

## References

1. Davies, P. J. The Plant Hormones: Their Nature, Occurrence, and Functions (Springer Netherlands, Dordrecht, 2010).

2. Santner, A., Calderon-Villalobos, L. I. A. & Estelle, M. Plant hormones are versatile chemical regulators of plant growth. Nat. Chem. Biol. 5, 301–307 (2009).

3. Tan, X. et al. Mechanism of auxin perception by the TIR1 ubiquitin ligase. Nature 446, 640 (2007).

4. Murase, K., Hirano, Y., Sun, T.-p. & Hakoshima, T. Gibberellin-induced DELLA recognition by the gibberellin receptor GID1. Nature 456, 459 (2008).

5. Sheard, L. B. et al. Jasmonate perception by inositol phosphate-potentiated COI1-JAZ co-receptor. Nature 468, 400 (2010).

6. Melcher, K. et al. A gate–latch–lock mechanism for hormone signalling by abscisic acid receptors. Nature 462, 602–608 (2009).

7. Miyazono, K.-i. et al. Structural basis of abscisic acid signalling. Nature 462, 609 (2009).

8. She, J. et al. Structural insight into brassinosteroid perception by BRI1. Nature 474, 472–476 (2011).

9. Hothorn, M. et al. Structural basis of steroid hormone perception by the receptor kinase BRI1. Nature 474, 467–471 (2011).

10. Hothorn, M., Dabi, T. & Chory, J. Structural basis for cytokinin recognition by *Arabidopsis thaliana* histidine kinase 4. Nat. Chem. Bio. 7, 766–768 (2011).

11. Kagiyama, M. et al. Structures of d14 and d14l in the strigolactone and karrikin signaling pathways. Genes Cells 18, 147–160 (2013).

12. Zhao, L.-H. et al. Destabilization of strigolactone receptor DWARF14 by binding of ligand ande3-ligase signaling effector DWARF3. Cell Res. 25, 1219–1236 (2015).

13. Wang, W. et al. Structural basis of salicylic acid perception by arabidopsis NPR proteins. Nature 586, 311–316 (2020).

14. Park, S.-Y. et al. Abscisic acid inhibits type 2c protein phosphatases via the PYR/PYL family of START proteins. Science (2009).

15. Ma, Y. et al. Regulators of PP2c phosphatase activity function as abscisic acid sensors. Science (2009).

16. Kobayashi, Y., Yamamoto, S., Minami, H., Kagaya, Y. & Hattori, T. Differential activation of the rice sucrose nonfermenting1—related protein kinase2 family by hyperosmotic stress and abscisic acid[w]. Plant Cell 16, 1163–1177 (2004).

17. Rigal, A., Ma, Q. & Robert, S. Unraveling plant hormone signaling through the use of small molecules. Front. Plant Sci. 5(2014).

18. Santiago, J., Henzler, C. & Hothorn, M. Molecular mechanism for plant steroid receptor activation by somatic embryogenesis co-receptor kinases. Science 341, 889–892 (2013).

19. Okamoto, M. et al. Activation of dimeric ABA receptors elicits guard cell closure, ABA-regulated gene expression, and drought tolerance. Proc. Natl. Acad. Sci. U.S.A. 110, 12132–12137 (2013).

20. Du, X. et al. Insights into protein—ligand interactions: Mechanisms, models, and methods. Int. J. Mol. Sci. 17, 144 (2016).

21. Breiten, B. et al. Water networks contribute to enthalpy/entropy compensation in pro-tein—ligand binding. J. Am. Chem. Soc. 135, 15579–15584 (2013).

22. Schiebel, J. et al. Intriguing role of water in protein-ligand binding studied by neutron crystallography on trypsin complexes. Nat. Commun. 9(2018).

23. Darby, J. F. et al. Water networks can determine the affinity of ligand binding to proteins. J. Am. Chem. Soc. 141, 15818–15826 (2019).

24. Haider, K., Wickstrom, L., Ramsey, S., Gilson, M. K. & Kurtzman, T. Enthalpic breakdown of water structure on protein active-site surfaces. J. Phys. Chem. B 120, 8743–8756 (2016).

25. Zsidó, B. Z. & Hetényi, C. The role of water in ligand binding. Curr Opin. Struct. Biol. 67, 1–8 (2021).

26. Hollingsworth, S. A. & Dror, R. O. Molecular dynamics simulation for all. Neuron 99, 1129–1143 (2018).

27. Buch, I., Giorgino, T. & Fabritiis, G. D. Complete reconstruction of an enzyme-inhibitor binding process by molecular dynamics simulations. Proc. Natl. Acad. Sci. U.S.A. 108, 10184–10189 (2011).

28. Lawrenz, M., Shukla, D. & Pande, V. S. Cloud computing approaches for prediction of ligand binding poses and pathways. Sci. Rep. 5, 7918 (2015).

29. Shukla, S., Zhao, C. & Shukla, D. Dewetting controls plant hormone perception and initiation of drought resistance signaling. Structure 27, 692–702.e3 (2019).

30. Aldukhi, F., Deb, A., Zhao, C., Moffett, A. S. & Shukla, D. Molecular mechanism of brassinosteroid perception by the plant growth receptor BRI1. J. Phys. Chem. B 124, 355–365 (2019).

31. Haider, K., Cruz, A., Ramsey, S., Gilson, M. K. & Kurtzman, T. Solvation structure and thermodynamic mapping (SSTMap): An open-source, flexible package for the analysis of water in molecular dynamics trajectories. J. Chem. Theory Comput. 14, 418–425 (2017).

32. Young, T., Abel, R., Kim, B., Berne, B. J. & Friesner, R. A. Motifs for molecular recognition exploiting hydrophobic enclosure in protein-ligand binding. Proc. Natl. Acad. Sci. U.S.A. 104, 808–813 (2007).

33. Abel, R., Young, T., Farid, R., Berne, B. J. & Friesner, R. A. Role of the active-site solvent in the thermodynamics of factor xa ligand binding. J. Am. Chem. Soc. 130, 2817–2831 (2008).

34. Chen, J., White, A., Nelson, D. C. & Shukla, D. Role of substrate recognition in modulating strigolactone receptor selectivity in witchweed. J. Biol. Chem. 101092 (2021).

35. Zhao, C. & Shukla, D. Molecular basis of the activation and dissociation of dimeric pyl2 receptor in abscisic acid signaling. bioRxiv 721761 (2021).

36. Moffett, A. S. & Shukla, D. Using molecular simulation to explore the nanoscale dynamics of the plant kinome. Biochem. J. 475, 905–921 (2018).

37. Feng, J., Chen, J., Selvam, B. & Shukla, D. Computational microscopy: Revealing molecular mechanisms in plants using molecular dynamics simulations. Plant Cell 31(2019).

38. Shukla, D., Hernández, C. X., Weber, J. K. & Pande, V. S. Markov state models provide insights into dynamic modulation of protein function. Acc. Chem. Res. 48, 414–422 (2015).

39. Husic, B. E. & Pande, V. S. Markov state models: From an art to a science. J. Am. Chem. Soc. 140, 2386–2396 (2018).

40. Soon, F.-F. et al. Abscisic acid signaling: Thermal stability shift assays as tool to analyze hormone perception and signal transduction. PLoS ONE 7, e47857 (2012).

41. Yuan, X. et al. Single amino acid alteration between valine and isoleucine determines the distinct pyrabactin selectivity by PYL1 and PYL2. J. Biol. Chem. 285, 28953–28958 (2010).

42. Balius, T. E. et al. Testing inhomogeneous solvation theory in structure-based ligand discovery. Proc. Natl. Acad. Sci. U. S. A. 114, E6839–E6846 (2017).

