## Supplementary Methods, Images, Tables and Results for "Intriguing Role of Water in Plant Hormone Perception"

### Electronic Supplementary Information (ESI)

#### Molecular Dynamics Simulations of the Binding of Plant Hormones to Receptors

**Initial structures**. Table S1 summarizes the details of the initial structures of plant hormones and their receptors, including TIR1/auxin (IAA), COI1/jasmonic acid (JA-Ile), GID1/gibberellin (GA3), AHK4/cytokinin (trans-zeatin), and NPR4/salicylic acid (SA) that were used for protein-ligand binding MD simulations. The protein and ligand structures were obtained from their crystal structures available in Protein Data Bank (PDB). For IAA and JA-Ile, their TIR1-ASK and COI1-ASK receptor complexes were used for MD simulations. Missing residues of COI1 were added by using the SWISS-MODEL webserver.[1] In both TIR1 and COI1, the leucine-rich repeat (LRR) domain contains an inositol phosphate cofactor ($InsP_6$ in TIR1 and $InsP_5$ in COI1). Since $InsP_5$ was not fully solved in the crystal structure of COI1, we drew an $InsP_5$ molecule and aligned $InsP_5$ to the structure of partially solved phosphates in 3OGK to obtain the initial structure of COI1-ASK complex. For salicylic acid (SA), the SA-binding core (SBC) of NPR4 was used for the simulations. Missing residues of the NPR4 SBC were modelled using the MODELLER package[2] (version 9.25) and the best structure was chosen based on its DOPE score.[3]

**Set up of MD simulations**. In order to study the binding of phytohormones to their receptors, we performed long timescale MD simulations to capture protein-ligand binding processes. Initially, we used Packmol[4] to randomly place the phytohormone molecules far away (at least 25 Å) from the binding sites of the plant hormone receptors. The receptor-ligand (also including cofactors $InsP_6$ in TIR1 and $InsP_5$ in COI1) complexes were then solvated with TIP3P water molecules to mimic solution environment. Certain number of $Na^+$ and $Cl^-$ ions were added to the system to neutralize the system and meet 150 mM salt concentrations. The Amber ff14SB force field was used for the proteins and the general Amber force field (GAFF)[5] was used for phytohormone molecules and cofactors ($InsP_5$ or $InsP_6$). The partial charges of phytohormone molecules were derived from Antechamber software using AM1-BCC method, and the partial charges of $InsP_5$ and $InsP_6$ were approximated from the previous theoretical study.[6] The simulation systems were minimized for 10,000 steps and then subjected to a series of heating steps to slowly increase the temperature of these systems to 300 K. Finally, the systems were equilibrated in isothermal-isobaric ensemble (300 K, 1 atm) for 1 ns. The simulations were run in isothermal-isobaric (300 K, 1 atm) ensemble using an integration time step of 2 fs. Periodic boundary condition was applied in all MD simulations. The particle-mesh Ewald method was used to treat the electrostatic interactions, along with a 10 Å cutoff distance for van der Waals interactions.[7] The SHAKE algorithm[8] was applied to constrain the length of covalent bonds involving hydrogen atoms.

**Steered MD simulations of GID1 receptor to open up N-terminal helix.** The crystal structure of GA3 receptor GID1 was in the active form, where the N-terminal helix closes the binding pocket at top of the binding site and interacts with GA3. In order to obtain the open conformation of GID1 without GA3 bound, we have performed 10 $\mu$s accelerated MD simulations to sample the conformational dynamics of GID1 in the absence of GA3 in the binding site. The simulations were performed in Amber 14 with steered MD simulation. We then clustered the conformations according to the N-terminal helix conformation, and selected a series of GID1 conformations to use as the starting structures for unbiased adaptive sampling MD simulations of protein-ligand binding.

**Accelerated MD simulations of NPR4 receptor to open up N-terminal helix.** The crystal structure of SA receptor (NPR4) was in its bound conformation where the N-terminal helix closes the binding pocket on top of the binding site. In order to obtain the open conformation of NPR4 without a bound ligand, we have performed 2 $\mu$s accelerated MD simulations to sample the conformational dynamics of NPR4 until SA escapes the binding pocket. The simulations were performed in Amber 18 employing the accelerated MD simulation method. Total and dihedral biasing terms were chosen following recommendations from Pierce *et al.*[9] We then clustered the conformations according to the protein conformation and ligand distance to the binding pocket, and selected a series of starting structures for unbiased adaptive sampling MD simulations of protein-ligand binding.

**Unbiased adaptive sampling of the binding of plant hormones to their receptors.** We used adaptive sampling to efficiently sample the protein-ligand binding and the conformational changes in the receptors. In the first round, the phytohormone molecules were placed far away from their receptors at multiple random configurations. MD simulations were launched from the equilibrated system and run for 70-100 ns. Then, a certain number of snapshots with the lowest protein-ligand distances were selected to serve as the starting structures for the next rounds of simulations. In the next round, short amount of MD simulations were performed starting from these configurations, with the initial velocities randomly assigned according to Boltzmann distribution. These iterative samplings were performed for multiple rounds and stopped until we have captured the protein-ligand binding and had enough simulation data to construct statistical models to describe the thermodynamics and kinetics of protein-ligand binding. Tables S2-S6 summarize the details of adaptive sampling for the IAA, JA-Ile, zeatin, GA3, and SA binding to their receptors, respectively.

### Markov State Model Construction and Hyperparameter Selection

**Featurization** Markov state model (MSM) is a powerful analytic technique to investigate large-scale unbiased MD simulation data on protein dynamics. MSMs discretize the protein conformational space into a certain number of individual conformational states and estimate the transition probabilities between these states, thereby allowing for investigation of protein dynamics at the timescale inaccessible via any conventional MD simulations. To discretize the protein conformational space, all the protein-ligand conformations collected from MD simulations were characterized by calculating a set of features that can best differentiate these configurations (featurization). In this work, we have calculated a set of distances between the atoms in protein and ligand to illustrate the protein-ligand interactions. Specifically, the CA atoms of the amino acids at the binding site (with the closest heavy-atom distance from the ligand within 5 Å) and heavy atoms in the phytohormones were calculated. For GID1, the residue-residue contacts between the binding site and the N-terminal helix were also calculated to characterize the conformational changes in GID1. For the AHK4, the residue-residue contacts between the binding site and the entry loop were also calculated to characterize the conformational changes in AHK4. For NPR4, the residue-residue distances between all $\alpha$-helices were used to capture the changes in protein conformation.

**Model construction and hyperparameter selection** Dimensionality reduction was then performed for clustering the conformation ensemble from MD simulations into kinetically relevant states. Time-lagged independent component analysis was used to identify several slowest degrees of freedom (tICs) resulted from linear combination of the original metrics.[10] K-means clusterings were then performed on the identified slowest motions to cluster the CG configurations into a certain number of states. MSMs were then constructed based on the clustering results. The matrix of transition probabilities was determined with a lag time $\tau$ using maximum likelihood approximation. The optimal $\tau$ was chosen based on the convergence of the implied timescales of MSMs and the number of clusters as well as the number of tICs were optimized via cross validation ranked by the variational GMRQ objective function (Figure S3).[11] All MSMs were constructed using the MSMBuilder 3.4[12] and the MSM hyperparameters were optimized using the Osprey software.[13] The MD simulation data for BLD binding to BRI1 (58 $\mu$s)[14], GR24 binding to D14 (198 $\mu$s)[15], and ABA binding to PYL2 (107 $\mu$s)[16] were taken from the past studies. The final MSM hyperparameters for the eight protein-ligand systems were summarized in Table S15.

### Transition Path Theory Analysis on Plant Hormone Binding Pathways

Transition path theory (TPT) was applied to calculate the probabilities and fluxes for the pathways between ligand-unbound and ligand-bound states in the protein-ligand binding MSMs.[17,18] The ligand-unbound and the ligand-bound states were chosen according to the distance between protein and ligand based on arbitrarily chosen cutoff values. Given the ligand-unbound and the ligand-bound MSM states for TPT analysis, numerous pathways that connected through the ligand-unbound states and the ligand-bound states were obtained, corresponding to plant hormone binding pathways. We reported the top pathways for these plant hormones with the greatest fluxes. The transition pathways were estimated using the MSMBuilder 3.4 python package.[12]

### Molecular Dynamics Simulations and Hydration Site Analysis of Plant Receptor Solvation Structure and Thermodynamic Properties

Explicit-solvent MD simulations on the *apo* and the *holo* plant hormone receptors, TIR1 (IAA), COI1 (JA-Ile), GID1 (GA3), AHK4 (zeatin), BRI1 (brassinolide), D14 (GR24), NPR4 (SA), and PYL2 (ABA) were performed to characterize the solvation structural and thermodynamic properties of their binding sites using Amber 18. The initial structures used for *apo* MD simulations were summarized in Table S1, and the initial structures used for the *holo* MD simulations were obtained from the binding MD simulations (except for NPR4, where the *holo* SBC crystal structure was available). The receptors were solvated with TIP3P water molecules, and $Na^+$ and $Cl^-$ ions were added to the system to neutralize the system and meet 150 mM salt concentrations. The systems were minimized for 20,000 steps and further equilibrated in NPT ensemble (1 atm, 300 K) for 1 ns to adjust the sizes of simulation boxes. To sample water densities for the binding sites efficiently, a recently developed MC/MD method[19] was used to equilibrate the system in isothermal-isovolumetric (NVT, 300 K) ensemble for 2500 MC/MD cycles. This methods allows the sampling of hydration sites even if they are buried in the protein. Each cycle includes 10000 MC steps that allow for water exchange between binding site and bulk as well as 1000 MD steps that relax water molecules in binding site. All heavy atoms of the receptor proteins were harmonically restrained using a force constant of 100 kcal·mol$^{-1}$·Å$^{-2}$. Production MD simulations were launched from the equilibrated structure and run for 100 ns in NVT with restraints applied to protein backbone using a force constant of 2.5 kcal·mol$^{-1}$·Å$^{-2}$. NVT was used in those production simulations to allow for faster convergence of sampled solvent density.[20] The coordinates of MD system were saved every 1 ps, resulting in trajectories with 100,000 frames. Then, the analysis calculations were performed using SSTMap python software (run_hsa program in SSTMap).[20] SSTMap calculates the structural and thermodynamic properties of water molecules on the surface of receptor cavity that are represented as a set of high-density hydration sites[21]. Briefly, the hydration sites were identified by clustering solvent distribution in the binding site into high-occupancy 1 Å spheres. Inhomogeneous solvation theory[22] was then used to compute the average system interaction energy and excess entropy terms for water in these hydration sites.

**Table S1.** Overview of the computational simulations and analysis method performed in this study. For receptor-ligand binding simulations, multiple parallel simulations were performed and their number of atoms and box sizes varied. The numbers provided in this table are average system size.

| **simulation** | **starting structure (PDB ID)** | **method** | **force field** | **software** | $\#_{atoms}$ | **size (Å$^3$)** | **ensemble** | **simulation time** |
|---|---|---|---|---|---|---|---|---|
| TIR1-ASK/IAA binding | 2P1Q | adaptive sampling, MD, MSM | Amber ff14SB, GAFF | Amber 14 | ~88,000 | ~95*106*103 | NPT | ~27.4 $\mu$s |
| COI1-ASK/JA-Ile binding | 3OGK | adaptive sampling, MD, MSM | Amber ff14SB, GAFF | Amber 14 | ~91,000 | ~93*114*103 | NPT | ~63.2 $\mu$s |
| GID1/GA3 binding | 2ZSH | adaptive sampling, MD, MSM | Amber ff14SB, GAFF | Amber 14 | ~85,000 | ~109*95*100 | NPT | ~129.5 $\mu$s |
| AHK4/Zeatin binding | 3T4L | adaptive sampling, MD, MSM | Amber ff14SB, GAFF | Amber 14 | ~80,000 | ~78*107*91 | NPT | ~36.7 $\mu$s |
| NPR4/SA binding | 6WPG | adaptive sampling, MD, MSM | Amber ff14SB, GAFF | Amber 18 | ~50,000 | ~90*90*90 | NPT | ~224.0 $\mu$s |
| TIR1 solvation | 2P1Q | MD, hydration site analysis | Amber ff14SB | Amber 18 | 84,725 | 93*107*102 | NVT | 100*2 ns |
| COI1 solvation | 3OGK | MD, hydration site analysis | Amber ff14SB | Amber 18 | 90,751 | 93*114*103 | NVT | 100*2 ns |
| GID1 solvation | 2ZSH | MD, hydration site analysis | Amber ff14SB | Amber 18 | 43,796 | 85*72*88 | NVT | 100*2 ns |
| D14 solvation | 4IH4 | MD, hydration site analysis | Amber ff14SB | Amber 18 | 32,233 | 74*73*75 | NVT | 100*2 ns |
| AHK4 solvation | 3T4L | MD, hydration site analysis | Amber ff14SB | Amber 18 | 42,701 | 69*72*108 | NVT | 100*2 ns |
| BRI1 solvation | 3RGZ | MD, hydration site analysis | Amber ff14SB | Amber 18 | 52,263 | 117*83*67 | NVT | 100*2 ns |
| NPR4 solvation | 6WPG | MD, hydration site analysis | Amber ff14SB | Amber 18 | 29,002 | 68*71*61 | NVT | 100*2 ns |
| PYL2 solvation | 3KDH | MD, hydration site analysis | Amber ff14SB | Amber 18 | 38,147 | 72*75*72 | NVT | 100*2 ns |

**Table S2.** Summary of the adaptive MD simulations of the binding of auxin to TIR1.

| Round | Parallel simulations | Simulation time (ns) | Aggregate ($\mu$s) |
|---|---|---|---|
| 1 | 100 | 80 | 8 |
| 2 | 100 | 100 | 10 |
| 3 | 120 | 60 | 7.2 |
| 4 | 40 | 60 | 2.4 |
| Total simulation time: ~27.4 $\mu$s | | | |

**Table S3.** Summary of the adaptive MD simulations of the binding of jasmonic acid to COI1.

| MD simulations starting from the snapshots where the ligand is far away from the binding site | | | |
|---|---|---|---|
| Round | Parallel simulations | Simulation time (ns) | Aggregate ($\mu$s) |
| 1 | 100 | 100 | 10 |
| 2 | 100 | 100 | 10 |
| 3 | 100 | 100 | 10 |
| 4 | 50 | 100 | 5 |
| 5 | 50 | 100 | 5 |
| 6 | 50 | 100 | 5 |
| 7 | 50 | 100 | 5 |
| 8 | 50 | 80 | 4 |
| 9 | 50 | 80 | 4 |
| MD simulations starting from the crystal structure where the ligand is bound to COI1 | | | |
| 1 | 20 | 100 | 2 |
| 2 | 50 | 100 | 5 |
| Total simulation time: ~63.2 $\mu$s | | | |

**Table S4.** Summary of the adaptive MD simulations of the binding of cytokinin to AHK4.

| Round | Parallel simulations | Simulation time (ns) | Aggregate ($\mu$s) |
|---|---|---|---|
| 1 | 100 | 76 | 7.6 |
| 2 | 50 | 120 | 7 |
| 3 | 100 | 120 | 12 |
| 4 | 100 | 120 | 12 |
| Total simulation time: ~36.7 $\mu$s | | | |

**Table S5.** Summary of the adaptive MD simulations of the binding of giberellin to GID1.

| Round | Parallel simulations | Simulation time (ns) | Aggregate ($\mu$s) |
|---|---|---|---|
| 1 | 200 | 78 | 15.6 |
| 2 | 60 | 112 | 6.7 |
| 3 | 100 | 112 | 11.2 |
| 4 | 50 | 112 | 5.6 |
| 5 | 100 | 84 | 8.4 |
| 6 | 100 | 112 | 11.2 |
| 7 | 50 | 112 | 5.6 |
| 8 | 100 | 84 | 8.4 |
| 9 | 600 | 100 | 60 |
| Total simulation time: ~129.5 $\mu$s | | | |

**Table S6.** Summary of the adaptive MD simulations of the binding of salicylic acid to NPR4.

| MD simulations starting from the crystal structure where the ligand is bound to NPR4 | | | |
|---|---|---|---|
| Round | Parallel simulations | Simulation time (ns) | Aggregate ($\mu$s) |
| 1 | 520 | 160 | 83 |
| MD simulations starting from the snapshots where the ligand is far away from the binding site | | | |
| 1 | 539 | 160 | 86 |
| 2 | 189 | 80 | 15 |
| MD simulations starting from the snapshots obtained by aMD | | | |
| 1 | 500 | 80 | 40 |
| Total simulation time: ~224 $\mu$s | | | |

**Table S7.** Hyperparameters for the MSMs constructed for the receptor-ligand binding simulations.

| **simulation** | **# tICs** | **lag time (ns)** | **# clusters** | **# MSM states** | **data utilization (%)** |
|---|---|---|---|---|---|
| TIR1/IAA binding | 6 | 30 | 100 | 100 | 100 |
| COI1/JA-Ile binding | 3 | 30 | 100 | 95 | 99.2 |
| GID1/GA3 binding | 2 | 30 | 200 | 200 | 100 |
| AHK4/Zeatin binding | 2 | 30 | 200 | 200 | 100 |
| BRI1/BLD binding | 6 | 20 | 200 | 200 | 100 |
| D14/GR24 binding | 4 | 20 | 325 | 325 | 100 |
| NPR4/SAL binding | 10 | 20 | 500 | 500 | 100 |
| PYL2/ABA binding | 4 | 30 | 300 | 300 | 100 |

**Table S8.** Calculated thermodynamic data (kcal/mol) and structural quantities for each of the 16 hydration sites in *apo* TIR1 receptor identified by clustering the active-site solvent density distribution. 'Neat' represents the pure TIP3P water.

| index | type | $f_o$ | $E_{sw}$ | $E_{ww}$ | $E_{tot}$ | $E_{ww}^{nbr}$ | $-TS^e$ | $N_{nbr}$ | $f_{enc}$ | $N_{sw}^{HB}$ | $N_{ww}^{HB}$ | $f_{ww}^{HB}$ | $N_{ww,lost}^{HB}$ |
|---|---|---|---|---|---|---|---|---|---|---|---|---|---|
| neat | - | - | 0.00 | -9.53 | -9.53 | -1.36 | 0.00 | 5.26 | 0.00 | 0.00 | 3.33 | 0.63 | 0.00 |
| 0 | P.Fr.U | 0.88 | -5.64 | -3.48 | -9.12 | -1.32 | 5.49 | 2.24 | 0.57 | 1.40 | 1.38 | 0.61 | 1.95 |
| 1 | C.Fr.F | 0.87 | -10.74 | -1.17 | -11.91 | -1.25 | 5.91 | 2.34 | 0.56 | 1.80 | 1.57 | 0.67 | 1.76 |
| 2 | P.En.F | 0.75 | -5.92 | -4.17 | -10.08 | -1.40 | 4.86 | 2.49 | 0.53 | 1.77 | 1.61 | 0.65 | 1.72 |
| 3 | P.En.F | 0.65 | -6.03 | -4.66 | -10.68 | -1.79 | 5.50 | 2.33 | 0.56 | 1.58 | 1.78 | 0.76 | 1.55 |
| 4 | C.Fr.F | 0.59 | -11.42 | 1.05 | -10.37 | 0.22 | 5.90 | 0.83 | 0.84 | 2.70 | 0.33 | 0.39 | 3.00 |
| 5 | P.En.F | 0.59 | -3.41 | -6.82 | -10.23 | -1.70 | 4.71 | 4.01 | 0.24 | 0.25 | 2.90 | 0.72 | 0.43 |
| 6 | C.Fr.U | 0.56 | -7.85 | -1.37 | -9.22 | -0.56 | 4.42 | 3.75 | 0.29 | 0.88 | 1.92 | 0.51 | 1.41 |
| 7 | C.Fr.F | 0.48 | -6.89 | -4.24 | -11.13 | -1.16 | 4.33 | 4.22 | 0.20 | 0.75 | 2.55 | 0.61 | 0.78 |
| 8 | A.En.U | 0.48 | -1.96 | -6.98 | -8.95 | -1.71 | 4.28 | 3.49 | 0.34 | 0.01 | 2.52 | 0.72 | 0.81 |
| 9 | P.En.F | 0.43 | -1.73 | -8.11 | -9.84 | -1.69 | 4.15 | 3.79 | 0.28 | 0.11 | 2.80 | 0.74 | 0.53 |
| 10 | P.Fr.F | 0.42 | -5.05 | -5.18 | -10.24 | -1.23 | 4.16 | 3.14 | 0.40 | 1.33 | 1.93 | 0.61 | 1.40 |
| 11 | P.En.F | 0.38 | -3.23 | -7.00 | -10.23 | -1.66 | 4.26 | 3.53 | 0.33 | 0.31 | 2.69 | 0.76 | 0.64 |
| 12 | C.En.F | 0.35 | -6.91 | -4.76 | -11.67 | -1.43 | 4.39 | 2.69 | 0.49 | 1.52 | 1.95 | 0.73 | 1.38 |
| 13 | C.En.F | 0.31 | -2.44 | -7.20 | -9.64 | -1.45 | 3.85 | 4.09 | 0.22 | 0.14 | 2.83 | 0.69 | 0.50 |
| 14 | P.En.U | 0.29 | -1.73 | -7.69 | -9.41 | -1.71 | 3.84 | 3.60 | 0.32 | 0.14 | 2.78 | 0.77 | 0.55 |
| 15 | P.Fr.F | 0.30 | -4.18 | -5.60 | -9.78 | -1.32 | 4.07 | 3.55 | 0.32 | 0.50 | 2.33 | 0.66 | 1.00 |

**Table S9.** Calculated thermodynamic data for each of the 23 hydration sites in *apo* COI1 receptor identified by clustering the active-site solvent density distribution. 'Neat' represents the pure TIP3P water.

| index | type | $f_o$ | $E_{sw}$ | $E_{ww}$ | $E_{tot}$ | $E_{ww}^{nbr}$ | $-TS^e$ | $N_{nbr}$ | $f_{enc}$ | $N_{sw}^{HB}$ | $N_{ww}^{HB}$ | $f_{ww}^{HB}$ | $N_{ww,lost}^{HB}$ |
|---|---|---|---|---|---|---|---|---|---|---|---|---|---|
| neat | - | - | 0.00 | -9.53 | -9.53 | -1.36 | 0.00 | 5.26 | 0.00 | 0.00 | 3.33 | 0.63 | 0.00 |
| 0 | C.Fr.F | 0.97 | -11.42 | 0.32 | -11.09 | -0.21 | 6.06 | 1.35 | 0.74 | 1.89 | 0.68 | 0.51 | 2.65 |
| 1 | P.En.F | 0.92 | -8.36 | -2.45 | -10.80 | -1.71 | 5.70 | 1.57 | 0.70 | 1.78 | 1.16 | 0.74 | 2.17 |
| 2 | C.En.F | 0.79 | -4.61 | -4.94 | -9.54 | -2.04 | 5.34 | 2.37 | 0.55 | 0.79 | 2.09 | 0.88 | 1.24 |
| 3 | C.Fr.F | 0.76 | -6.54 | -3.08 | -9.63 | -1.13 | 4.91 | 2.54 | 0.52 | 1.86 | 1.50 | 0.59 | 1.83 |
| 4 | C.En.U | 0.72 | -4.85 | -4.32 | -9.17 | -2.04 | 5.07 | 1.21 | 0.77 | 1.81 | 0.94 | 0.78 | 2.39 |
| 5 | C.Fr.U | 0.7 | -6.61 | -2.50 | -9.11 | -1.26 | 4.61 | 2.13 | 0.60 | 0.88 | 1.40 | 0.66 | 1.93 |
| 6 | C.Fr.F | 0.61 | -6.96 | -4.75 | -11.71 | -1.27 | 4.86 | 3.50 | 0.33 | 1.01 | 2.35 | 0.67 | 0.98 |
| 7 | C.En.F | 0.62 | -6.28 | -4.69 | -10.97 | -1.51 | 4.86 | 2.21 | 0.58 | 1.45 | 1.61 | 0.73 | 1.72 |
| 8 | C.Fr.F | 0.59 | -8.44 | -3.21 | -11.65 | -1.01 | 4.80 | 3.40 | 0.35 | 1.45 | 2.01 | 0.59 | 1.32 |
| 9 | P.En.F | 0.55 | -7.86 | -2.85 | -10.71 | -1.59 | 4.88 | 2.45 | 0.54 | 1.44 | 1.80 | 0.74 | 1.53 |
| 10 | C.Fr.F | 0.55 | -9.58 | -1.60 | -11.18 | -0.77 | 4.59 | 3.74 | 0.29 | 0.33 | 1.97 | 0.53 | 1.36 |
| 11 | C.Fr.F | 0.53 | -8.14 | -3.26 | -11.39 | -1.01 | 4.63 | 3.86 | 0.27 | 1.21 | 2.26 | 0.59 | 1.07 |
| 12 | A.En.U | 0.47 | -2.47 | -6.59 | -9.06 | -2.08 | 4.56 | 3.17 | 0.40 | 0.02 | 2.64 | 0.83 | 0.69 |
| 13 | P.En.U | 0.42 | -2.68 | -6.75 | -9.44 | -1.59 | 3.98 | 3.79 | 0.28 | 0.25 | 2.75 | 0.73 | 0.58 |
| 14 | A.En.U | 0.41 | -3.70 | -5.79 | -9.49 | -1.67 | 4.62 | 3.34 | 0.37 | 0.00 | 2.59 | 0.78 | 0.74 |
| 15 | C.En.F | 0.39 | -6.24 | -3.96 | -10.20 | -1.73 | 4.24 | 2.28 | 0.57 | 0.92 | 1.82 | 0.80 | 1.51 |
| 16 | A.En.F | 0.36 | -3.90 | -7.21 | -11.11 | -1.61 | 4.54 | 4.29 | 0.18 | 0.02 | 3.34 | 0.78 | -0.01 |
| 17 | C.Fr.F | 0.35 | -7.89 | -3.23 | -11.12 | -1.17 | 3.96 | 2.71 | 0.48 | 1.81 | 1.70 | 0.63 | 1.63 |
| 18 | A.En.U | 0.33 | -2.42 | -6.17 | -8.60 | -1.79 | 4.01 | 2.94 | 0.44 | 0.02 | 2.22 | 0.76 | 1.11 |
| 19 | P.Fr.F | 0.32 | -6.75 | -3.59 | -10.34 | -0.97 | 4.15 | 3.82 | 0.27 | 1.14 | 2.16 | 0.57 | 1.17 |
| 20 | P.En.U | 0.32 | -4.81 | -2.72 | -7.53 | -1.73 | 3.89 | 1.52 | 0.71 | 0.79 | 1.18 | 0.78 | 2.15 |
| 21 | P.En.F | 0.3 | -5.26 | -4.60 | -9.86 | -1.72 | 4.58 | 2.41 | 0.54 | 0.50 | 1.85 | 0.77 | 1.48 |
| 22 | C.Fr.F | 0.3 | -8.69 | -1.70 | -10.39 | -1.10 | 4.22 | 1.73 | 0.67 | 1.59 | 1.14 | 0.66 | 2.19 |

**Table S10.** Calculated thermodynamic data for each of the 20 hydration sites in *apo* BRI1 receptor identified by clustering the active-site solvent density distribution. 'Neat' represents the pure TIP3P water.

| index | type | $f_o$ | $E_{sw}$ | $E_{ww}$ | $E_{tot}$ | $E_{ww}^{nbr}$ | $-TS^e$ | $N_{nbr}$ | $f_{enc}$ | $N_{sw}^{HB}$ | $N_{ww}^{HB}$ | $f_{ww}^{HB}$ | $N_{ww,lost}^{HB}$ |
|---|---|---|---|---|---|---|---|---|---|---|---|---|---|
| neat | - | - | 0.00 | -9.53 | -9.53 | -1.36 | 0.00 | 5.26 | 0.00 | 0.00 | 3.33 | 0.63 | 0.00 |
| 0 | C.En.F | 0.64 | -6.72 | -3.33 | -10.05 | -2.11 | 5.30 | 1.77 | 0.66 | 1.31 | 1.49 | 0.85 | 1.84 |
| 1 | P.En.U | 0.6 | -3.77 | -5.17 | -8.94 | -1.56 | 4.65 | 2.38 | 0.55 | 0.56 | 1.73 | 0.73 | 1.60 |
| 2 | C.En.U | 0.55 | -4.58 | -4.36 | -8.93 | -2.02 | 4.88 | 1.94 | 0.63 | 0.95 | 1.64 | 0.84 | 1.69 |
| 3 | P.Fr.U | 0.46 | -3.77 | -5.63 | -9.40 | -1.30 | 4.22 | 3.17 | 0.40 | 1.12 | 2.05 | 0.65 | 1.28 |
| 4 | P.En.U | 0.43 | -3.39 | -6.01 | -9.41 | -1.44 | 3.95 | 3.05 | 0.42 | 0.86 | 2.08 | 0.68 | 1.25 |
| 5 | P.En.F | 0.39 | -3.40 | -6.25 | -9.65 | -1.56 | 4.25 | 3.35 | 0.36 | 0.79 | 2.39 | 0.71 | 0.94 |
| 6 | A.En.U | 0.39 | -1.10 | -8.07 | -9.17 | -1.74 | 3.86 | 3.66 | 0.30 | 0.01 | 2.82 | 0.77 | 0.51 |
| 7 | P.En.F | 0.36 | -5.65 | -4.82 | -10.46 | -1.53 | 4.76 | 3.04 | 0.42 | 0.99 | 2.27 | 0.75 | 1.06 |
| 8 | P.En.F | 0.36 | -3.71 | -6.36 | -10.07 | -1.48 | 4.07 | 3.15 | 0.40 | 1.25 | 2.05 | 0.65 | 1.28 |
| 9 | P.En.U | 0.35 | -2.96 | -6.01 | -8.97 | -1.47 | 3.99 | 3.54 | 0.33 | 0.66 | 2.32 | 0.65 | 1.01 |
| 10 | P.Fr.U | 0.34 | -2.95 | -6.50 | -9.45 | -1.32 | 3.71 | 3.56 | 0.32 | 1.08 | 2.12 | 0.59 | 1.21 |
| 11 | A.En.U | 0.32 | -0.95 | -7.36 | -8.31 | -1.73 | 3.85 | 3.05 | 0.42 | 0.00 | 2.29 | 0.75 | 1.04 |
| 12 | P.En.U | 0.32 | -1.48 | -8.00 | -9.48 | -1.44 | 3.60 | 4.05 | 0.23 | 0.73 | 2.66 | 0.66 | 0.67 |
| 13 | P.En.U | 0.32 | -3.34 | -5.91 | -9.25 | -1.39 | 3.66 | 3.47 | 0.34 | 0.89 | 2.28 | 0.66 | 1.05 |
| 14 | P.En.F | 0.31 | -4.45 | -5.84 | -10.29 | -1.79 | 4.19 | 2.75 | 0.48 | 1.10 | 2.15 | 0.78 | 1.18 |
| 15 | P.Fr.F | 0.31 | -3.90 | -6.03 | -9.93 | -1.19 | 3.76 | 4.02 | 0.24 | 0.87 | 2.46 | 0.61 | 0.87 |
| 16 | C.Fr.U | 0.3 | -4.13 | -5.22 | -9.35 | -1.26 | 3.71 | 3.28 | 0.38 | 0.91 | 2.00 | 0.61 | 1.33 |
| 17 | A.En.U | 0.29 | -2.01 | -7.36 | -9.38 | -1.58 | 3.81 | 3.76 | 0.28 | 0.00 | 2.66 | 0.71 | 0.67 |
| 18 | P.En.U | 0.29 | -2.91 | -6.29 | -9.20 | -1.52 | 3.67 | 3.17 | 0.40 | 0.62 | 2.25 | 0.71 | 1.08 |
| 19 | A.En.U | 0.29 | -1.63 | -6.02 | -7.65 | -1.80 | 3.77 | 2.64 | 0.50 | 0.02 | 2.03 | 0.77 | 1.30 |

**Table S11.** Calculated thermodynamic data for each of the 29 hydration sites in *apo* D14 receptor identified by clustering the active-site solvent density distribution. 'Neat' represents the pure TIP3P water.

| index | type | $f_o$ | $E_{sw}$ | $E_{ww}$ | $E_{tot}$ | $E_{ww}^{nbr}$ | $-TS^e$ | $N_{nbr}$ | $f_{enc}$ | $N_{sw}^{HB}$ | $N_{ww}^{HB}$ | $f_{ww}^{HB}$ | $N_{ww,lost}^{HB}$ |
|---|---|---|---|---|---|---|---|---|---|---|---|---|---|
| neat | - | - | 0.00 | -9.53 | -9.53 | -1.36 | 0.00 | 5.26 | 0.00 | 0.00 | 3.33 | 0.63 | 0.00 |
| 0 | P.En.F | 0.85 | -3.60 | -5.98 | -9.57 | -1.82 | 4.90 | 2.79 | 0.47 | 0.77 | 2.10 | 0.75 | 1.23 |
| 1 | P.En.F | 0.84 | -7.04 | -2.72 | -9.75 | -1.85 | 5.50 | 1.30 | 0.75 | 1.19 | 1.06 | 0.81 | 2.27 |
| 2 | P.En.F | 0.73 | -4.91 | -4.70 | -9.62 | -1.79 | 5.10 | 2.40 | 0.54 | 0.46 | 1.99 | 0.83 | 1.34 |
| 3 | C.Fr.F | 0.67 | -7.27 | -4.30 | -11.58 | -0.99 | 4.96 | 3.42 | 0.35 | 0.96 | 2.06 | 0.60 | 1.27 |
| 4 | P.En.U | 0.69 | -3.77 | -4.59 | -8.35 | -2.05 | 4.81 | 1.95 | 0.63 | 0.55 | 1.71 | 0.88 | 1.62 |
| 5 | P.En.U | 0.65 | -5.37 | -3.07 | -8.44 | -2.04 | 5.14 | 1.47 | 0.72 | 0.74 | 1.24 | 0.84 | 2.09 |
| 6 | A.En.F | 0.64 | -2.87 | -7.34 | -10.20 | -1.62 | 5.46 | 3.47 | 0.34 | 0.00 | 2.72 | 0.78 | 0.61 |
| 7 | P.En.U | 0.53 | -1.88 | -6.88 | -8.75 | -1.44 | 4.30 | 3.50 | 0.33 | 0.32 | 2.41 | 0.69 | 0.92 |
| 8 | P.En.F | 0.52 | -3.88 | -5.86 | -9.75 | -1.73 | 4.79 | 3.17 | 0.40 | 0.51 | 2.44 | 0.77 | 0.89 |
| 9 | C.Fr.F | 0.47 | -5.38 | -4.51 | -9.89 | -1.33 | 4.78 | 3.11 | 0.41 | 0.76 | 1.89 | 0.61 | 1.44 |
| 10 | P.En.U | 0.46 | -4.45 | -4.82 | -9.27 | -1.59 | 4.41 | 2.36 | 0.55 | 0.95 | 1.79 | 0.76 | 1.54 |
| 11 | A.En.U | 0.42 | -3.14 | -5.68 | -8.83 | -1.77 | 4.51 | 2.76 | 0.48 | 0.00 | 2.11 | 0.77 | 1.22 |
| 12 | A.En.F | 0.42 | -3.59 | -7.32 | -10.91 | -1.57 | 4.49 | 4.29 | 0.19 | 0.00 | 3.35 | 0.78 | -0.02 |
| 13 | A.En.F | 0.42 | -1.66 | -8.68 | -10.34 | -1.61 | 4.66 | 4.57 | 0.13 | 0.00 | 3.34 | 0.73 | -0.01 |
| 14 | C.En.F | 0.4 | -7.44 | -3.73 | -11.17 | -1.51 | 4.93 | 2.51 | 0.52 | 1.60 | 1.79 | 0.72 | 1.54 |
| 15 | A.En.U | 0.4 | -1.19 | -7.81 | -9.00 | -1.69 | 3.98 | 3.64 | 0.31 | 0.00 | 2.82 | 0.77 | 0.51 |
| 16 | P.En.F | 0.38 | -4.08 | -5.89 | -9.97 | -1.52 | 4.35 | 2.94 | 0.44 | 0.66 | 2.15 | 0.73 | 1.18 |
| 17 | A.En.U | 0.37 | -1.53 | -7.45 | -8.97 | -1.72 | 4.25 | 3.46 | 0.34 | 0.00 | 2.65 | 0.77 | 0.68 |
| 18 | A.En.U | 0.37 | -0.78 | -8.26 | -9.04 | -1.56 | 4.02 | 3.97 | 0.24 | 0.00 | 2.97 | 0.75 | 0.36 |
| 19 | P.En.U | 0.36 | -1.59 | -7.70 | -9.29 | -1.65 | 4.21 | 3.57 | 0.32 | 0.24 | 2.74 | 0.77 | 0.59 |
| 20 | A.En.U | 0.36 | -3.39 | -6.14 | -9.53 | -1.61 | 4.18 | 3.46 | 0.34 | 0.01 | 2.69 | 0.78 | 0.64 |
| 21 | A.En.U | 0.35 | -1.09 | -8.34 | -9.43 | -1.62 | 4.12 | 4.10 | 0.22 | 0.04 | 3.00 | 0.73 | 0.33 |
| 22 | A.En.F | 0.35 | -2.97 | -6.83 | -9.80 | -1.72 | 4.57 | 3.76 | 0.28 | 0.00 | 2.95 | 0.78 | 0.38 |
| 23 | P.En.U | 0.33 | -3.26 | -4.78 | -8.05 | -1.44 | 3.64 | 2.89 | 0.45 | 0.32 | 1.98 | 0.68 | 1.35 |
| 24 | A.En.F | 0.33 | -1.58 | -8.18 | -9.75 | -1.64 | 3.76 | 4.29 | 0.18 | 0.01 | 3.16 | 0.74 | 0.17 |
| 25 | A.En.F | 0.31 | -2.27 | -7.79 | -10.06 | -1.83 | 4.57 | 3.75 | 0.29 | 0.00 | 3.08 | 0.82 | 0.25 |
| 26 | A.En.F | 0.31 | -1.69 | -7.98 | -9.67 | -1.44 | 3.84 | 4.64 | 0.12 | 0.01 | 3.22 | 0.69 | 0.11 |
| 27 | P.En.U | 0.3 | -2.80 | -6.58 | -9.37 | -1.50 | 3.75 | 3.49 | 0.34 | 0.88 | 2.39 | 0.68 | 0.94 |
| 28 | A.En.U | 0.3 | -0.80 | -8.30 | -9.10 | -1.53 | 3.67 | 4.35 | 0.17 | 0.09 | 2.94 | 0.68 | 0.39 |

**Table S12.** Calculated thermodynamic data for each of the 22 hydration sites in *apo* GID1 receptor identified by clustering the active-site solvent density distribution. 'Neat' represents the pure TIP3P water.

| index | type | $f_o$ | $E_{sw}$ | $E_{ww}$ | $E_{tot}$ | $E_{ww}^{nbr}$ | $-TS^e$ | $N_{nbr}$ | $f_{enc}$ | $N_{sw}^{HB}$ | $N_{ww}^{HB}$ | $f_{ww}^{HB}$ | $N_{ww,lost}^{HB}$ |
|---|---|---|---|---|---|---|---|---|---|---|---|---|---|
| neat | - | - | 0.00 | -9.53 | -9.53 | -1.36 | 0.00 | 5.26 | 0.00 | 0.00 | 3.33 | 0.63 | 0.00 |
| 0 | C.En.F | 0.9 | -6.23 | -4.02 | -10.25 | -1.37 | 5.61 | 2.59 | 0.51 | 1.43 | 1.82 | 0.70 | 1.51 |
| 1 | C.En.F | 0.89 | -9.57 | -2.69 | -12.26 | -1.41 | 6.14 | 2.64 | 0.50 | 1.91 | 1.72 | 0.65 | 1.61 |
| 2 | P.Fr.F | 0.85 | -9.28 | -1.42 | -10.69 | -0.75 | 5.42 | 2.26 | 0.57 | 1.03 | 1.45 | 0.64 | 1.88 |
| 3 | P.En.F | 0.74 | -6.29 | -4.13 | -10.42 | -1.55 | 5.92 | 2.21 | 0.58 | 1.10 | 1.86 | 0.84 | 1.47 |
| 4 | C.En.F | 0.73 | -6.13 | -5.12 | -11.24 | -1.65 | 5.30 | 3.14 | 0.40 | 1.18 | 2.60 | 0.83 | 0.73 |
| 5 | C.En.U | 0.69 | -2.94 | -6.37 | -9.31 | -1.70 | 4.80 | 3.03 | 0.42 | 0.71 | 2.19 | 0.72 | 1.14 |
| 6 | C.Fr.F | 0.68 | -8.10 | -3.39 | -11.49 | -1.12 | 5.21 | 3.13 | 0.41 | 1.19 | 2.06 | 0.66 | 1.27 |
| 7 | C.Fr.F | 0.64 | -7.37 | -2.27 | -9.64 | -1.13 | 5.25 | 2.49 | 0.53 | 0.92 | 1.54 | 0.62 | 1.79 |
| 8 | P.Fr.F | 0.63 | -5.67 | -3.96 | -9.63 | -1.21 | 4.63 | 2.48 | 0.53 | 1.05 | 1.71 | 0.69 | 1.62 |
| 9 | P.Fr.F | 0.62 | -6.27 | -4.06 | -10.33 | -1.17 | 4.72 | 2.92 | 0.45 | 1.26 | 1.83 | 0.63 | 1.50 |
| 10 | C.En.U | 0.6 | -2.33 | -6.16 | -8.49 | -2.39 | 4.86 | 2.08 | 0.60 | 0.63 | 1.92 | 0.92 | 1.41 |
| 11 | P.En.F | 0.54 | -6.15 | -4.15 | -10.30 | -1.66 | 4.94 | 1.98 | 0.62 | 1.83 | 1.40 | 0.71 | 1.93 |
| 12 | P.En.F | 0.48 | -2.68 | -7.05 | -9.73 | -1.64 | 4.19 | 3.69 | 0.30 | 0.12 | 2.83 | 0.77 | 0.50 |
| 13 | P.En.F | 0.47 | -3.36 | -6.65 | -10.01 | -1.55 | 4.83 | 3.36 | 0.36 | 0.86 | 2.68 | 0.80 | 0.65 |
| 14 | P.Fr.U | 0.4 | -4.41 | -5.03 | -9.44 | -1.20 | 4.29 | 3.99 | 0.24 | 0.65 | 2.48 | 0.62 | 0.85 |
| 15 | P.En.F | 0.39 | -5.16 | -4.89 | -10.05 | -1.38 | 4.50 | 3.72 | 0.29 | 0.47 | 2.54 | 0.68 | 0.79 |
| 16 | P.En.U | 0.37 | -0.91 | -6.91 | -7.81 | -1.87 | 4.27 | 2.65 | 0.50 | 0.13 | 1.96 | 0.74 | 1.37 |
| 17 | A.En.U | 0.34 | -1.63 | -7.89 | -9.52 | -1.96 | 4.16 | 3.31 | 0.37 | 0.02 | 2.73 | 0.83 | 0.60 |
| 18 | A.En.U | 0.32 | -3.34 | -5.82 | -9.17 | -1.51 | 4.19 | 3.70 | 0.30 | 0.01 | 2.60 | 0.70 | 0.73 |
| 19 | P.En.U | 0.3 | -2.91 | -5.53 | -8.44 | -1.53 | 4.00 | 2.90 | 0.45 | 0.52 | 2.01 | 0.69 | 1.32 |
| 20 | P.En.F | 0.3 | -6.38 | -3.21 | -9.59 | -1.66 | 4.15 | 1.64 | 0.69 | 1.96 | 1.30 | 0.80 | 2.03 |
| 21 | P.Fr.F | 0.3 | -8.87 | -3.91 | -12.78 | -1.36 | 5.25 | 2.96 | 0.44 | 1.61 | 2.13 | 0.72 | 1.20 |

**Table S13.** Calculated thermodynamic data for each of the 15 hydration sites in *apo* AHK4 receptor identified by clustering the active-site solvent density distribution. 'Neat' represents the pure TIP3P water.

| index | type | $f_o$ | $E_{sw}$ | $E_{ww}$ | $E_{tot}$ | $E_{ww}^{nbr}$ | $-TS^e$ | $N_{nbr}$ | $f_{enc}$ | $N_{sw}^{HB}$ | $N_{ww}^{HB}$ | $f_{ww}^{HB}$ | $N_{ww,lost}^{HB}$ |
|---|---|---|---|---|---|---|---|---|---|---|---|---|---|
| neat | - | - | 0.00 | -9.53 | -9.53 | -1.36 | 0.00 | 5.26 | 0.00 | 0.00 | 3.33 | 0.63 | 0.00 |
| 0 | P.En.F | 0.8 | -4.80 | -6.37 | -11.17 | -2.04 | 5.45 | 2.91 | 0.45 | 1.01 | 2.61 | 0.90 | 0.72 |
| 1 | C.En.U | 0.73 | -5.79 | -3.69 | -9.47 | -1.56 | 4.77 | 2.58 | 0.51 | 0.88 | 2.03 | 0.78 | 1.30 |
| 2 | P.En.F | 0.72 | -5.00 | -5.95 | -10.95 | -2.14 | 5.34 | 2.14 | 0.59 | 1.56 | 1.88 | 0.88 | 1.45 |
| 3 | C.Fr.U | 0.58 | -7.44 | -1.17 | -8.60 | -0.79 | 4.94 | 2.76 | 0.48 | 1.24 | 1.51 | 0.55 | 1.82 |
| 4 | A.En.U | 0.57 | -2.83 | -5.63 | -8.46 | -1.77 | 4.42 | 3.20 | 0.39 | 0.00 | 2.46 | 0.77 | 0.87 |
| 5 | P.En.U | 0.56 | -1.87 | -6.70 | -8.58 | -2.08 | 4.53 | 2.77 | 0.47 | 0.12 | 2.43 | 0.88 | 0.90 |
| 6 | C.Fr.F | 0.55 | -8.40 | -2.42 | -10.83 | -1.07 | 4.89 | 2.56 | 0.51 | 1.22 | 1.56 | 0.61 | 1.77 |
| 7 | P.En.U | 0.5 | -4.92 | -4.42 | -9.34 | -2.17 | 4.37 | 2.10 | 0.60 | 0.88 | 1.77 | 0.84 | 1.56 |
| 8 | P.En.U | 0.43 | -2.63 | -5.36 | -8.00 | -2.38 | 4.60 | 1.84 | 0.65 | 0.88 | 1.70 | 0.92 | 1.63 |
| 9 | C.En.U | 0.39 | -2.47 | -6.02 | -8.50 | -1.45 | 4.24 | 3.27 | 0.38 | 0.89 | 2.21 | 0.67 | 1.12 |
| 10 | C.Fr.F | 0.38 | -7.64 | -2.47 | -10.11 | -1.16 | 4.31 | 2.75 | 0.48 | 1.05 | 1.58 | 0.57 | 1.75 |
| 11 | P.Fr.U | 0.33 | -8.55 | -0.01 | -8.56 | -0.95 | 4.10 | 1.12 | 0.79 | 1.52 | 0.58 | 0.52 | 2.75 |
| 12 | P.En.U | 0.31 | -2.28 | -5.88 | -8.16 | -1.97 | 4.06 | 2.60 | 0.50 | 0.36 | 2.02 | 0.78 | 1.31 |
| 13 | A.Fr.U | 0.3 | 0.16 | -5.45 | -5.29 | -1.10 | 3.79 | 4.00 | 0.24 | 0.07 | 2.35 | 0.59 | 0.98 |
| 14 | A.En.U | 0.29 | -2.62 | -4.71 | -7.34 | -1.99 | 3.55 | 2.16 | 0.59 | 0.01 | 1.80 | 0.83 | 1.53 |

**Table S14.** Calculated thermodynamic data for each of the 4 hydration sites in *apo* NPR4 receptor identified by clustering the active-site solvent density distribution. 'Neat' represents the pure TIP3P water.

| index | type | $f_o$ | $E_{sw}$ | $E_{ww}$ | $E_{tot}$ | $E_{ww}^{nbr}$ | $-TS^e$ | $N_{nbr}$ | $f_{enc}$ | $N_{sw}^{HB}$ | $N_{ww}^{HB}$ | $f_{ww}^{HB}$ | $N_{ww,lost}^{HB}$ |
|---|---|---|---|---|---|---|---|---|---|---|---|---|---|
| neat | - | - | 0.00 | -9.53 | -9.53 | -1.36 | 0.00 | 5.26 | 0.00 | 0.00 | 3.33 | 0.63 | 0.00 |
| 0 | C.En.U | 0.97 | -6.23 | -2.14 | -8.37 | -2.32 | 4.52 | 0.90 | 0.83 | 1.05 | 0.62 | 0.62 | 2.78 |
| 1 | C.En.U | 0.62 | -4.43 | -2.14 | -6.57 | -2.37 | 3.99 | 0.93 | 0.82 | 0.48 | 0.78 | 0.78 | 2.74 |
| 2 | P.En.U | 0.59 | -3.46 | -4.27 | -7.73 | -2.34 | 4.33 | 1.86 | 0.65 | 0.32 | 1.39 | 0.75 | 2.15 |
| 3 | P.En.U | 0.41 | -3.92 | -3.80 | -7.72 | -2.32 | 4.57 | 1.54 | 0.70 | 0.74 | 1.22 | 0.82 | 2.36 |

**Table S15.** Calculated thermodynamic data for each of the 30 hydration sites in *apo* PYL2 receptor identified by clustering the active-site solvent density distribution. 'Neat' represents the pure TIP3P water.

| index | type | $f_o$ | $E_{sw}$ | $E_{ww}$ | $E_{tot}$ | $E_{ww}^{nbr}$ | $-TS^e$ | $N_{nbr}$ | $f_{enc}$ | $N_{sw}^{HB}$ | $N_{ww}^{HB}$ | $f_{ww}^{HB}$ | $N_{ww,lost}^{HB}$ |
|---|---|---|---|---|---|---|---|---|---|---|---|---|---|
| neat | - | - | 0.00 | -9.53 | -9.53 | -1.36 | 0.00 | 5.26 | 0.00 | 0.00 | 3.33 | 0.63 | 0.00 |
| 0 | C.Fr.F | 0.94 | -14.42 | 1.76 | -12.66 | -0.29 | 6.25 | 2.14 | 0.59 | 1.96 | 1.31 | 0.61 | 2.02 |
| 1 | C.Fr.F | 0.88 | -13.61 | 1.57 | -12.05 | -0.13 | 6.20 | 1.26 | 0.76 | 2.85 | 0.50 | 0.40 | 2.83 |
| 2 | C.En.U | 0.87 | -2.98 | -6.17 | -9.15 | -1.55 | 4.73 | 3.00 | 0.43 | 0.89 | 2.29 | 0.76 | 1.04 |
| 3 | C.Fr.F | 0.86 | -12.32 | 0.74 | -11.58 | -0.35 | 5.67 | 2.09 | 0.60 | 2.05 | 1.01 | 0.48 | 2.32 |
| 4 | C.Fr.F | 0.84 | -14.63 | 3.42 | -11.21 | 0.26 | 5.72 | 2.85 | 0.46 | 1.77 | 1.14 | 0.40 | 2.19 |
| 5 | C.Fr.F | 0.78 | -7.11 | -3.17 | -10.28 | -1.08 | 5.19 | 3.35 | 0.36 | 1.00 | 1.98 | 0.59 | 1.35 |
| 6 | C.Fr.F | 0.75 | -13.62 | 0.33 | -13.30 | -0.65 | 6.49 | 1.16 | 0.78 | 2.91 | 0.86 | 0.74 | 2.47 |
| 7 | C.En.F | 0.72 | -4.11 | -5.74 | -9.84 | -1.88 | 5.15 | 2.76 | 0.47 | 0.96 | 2.31 | 0.83 | 1.02 |
| 8 | A.En.U | 0.69 | -1.42 | -7.96 | -9.38 | -1.90 | 4.25 | 3.55 | 0.33 | 0.06 | 2.89 | 0.82 | 0.44 |
| 9 | C.Fr.F | 0.66 | -6.02 | -3.66 | -9.68 | -0.96 | 4.75 | 3.42 | 0.35 | 0.93 | 1.83 | 0.54 | 1.50 |
| 10 | C.Fr.F | 0.64 | -9.01 | -0.97 | -9.99 | -0.85 | 5.10 | 1.55 | 0.71 | 1.57 | 0.93 | 0.60 | 2.40 |
| 11 | C.En.U | 0.64 | -4.27 | -4.75 | -9.02 | -1.41 | 4.58 | 2.97 | 0.43 | 0.95 | 2.04 | 0.69 | 1.29 |
| 12 | A.En.U | 0.59 | -3.36 | -5.70 | -9.06 | -1.63 | 4.74 | 3.38 | 0.36 | 0.01 | 2.72 | 0.80 | 0.61 |
| 13 | P.En.U | 0.52 | -3.29 | -6.21 | -9.50 | -1.53 | 4.39 | 3.39 | 0.35 | 0.84 | 2.36 | 0.69 | 0.97 |
| 14 | P.En.F | 0.53 | -3.28 | -6.70 | -9.98 | -1.64 | 4.69 | 3.21 | 0.39 | 0.93 | 2.39 | 0.74 | 0.94 |
| 15 | A.En.U | 0.50 | -1.86 | -7.57 | -9.42 | -1.66 | 4.12 | 3.73 | 0.29 | 0.00 | 2.64 | 0.71 | 0.69 |
| 16 | P.En.U | 0.50 | -5.87 | -3.29 | -9.16 | -1.58 | 4.23 | 1.63 | 0.69 | 1.61 | 1.18 | 0.72 | 2.15 |
| 17 | C.Fr.U | 0.49 | -10.10 | 0.79 | -9.31 | -0.94 | 5.51 | 1.06 | 0.80 | 1.13 | 0.89 | 0.85 | 2.44 |
| 18 | A.En.F | 0.48 | -1.42 | -8.15 | -9.57 | -1.59 | 3.93 | 4.36 | 0.17 | 0.01 | 3.20 | 0.73 | 0.13 |
| 19 | A.En.F | 0.40 | -0.76 | -8.77 | -9.53 | -1.62 | 3.78 | 4.45 | 0.15 | 0.02 | 3.22 | 0.72 | 0.11 |
| 20 | C.En.U | 0.39 | -3.27 | -5.20 | -8.47 | -1.69 | 4.01 | 2.65 | 0.50 | 0.59 | 1.96 | 0.74 | 1.37 |
| 21 | C.Fr.F | 0.38 | -7.36 | -2.18 | -9.53 | -0.97 | 4.46 | 3.45 | 0.34 | 0.76 | 2.05 | 0.60 | 1.28 |
| 22 | A.En.F | 0.35 | -0.94 | -8.85 | -9.79 | -1.63 | 3.66 | 4.40 | 0.16 | 0.01 | 3.24 | 0.74 | 0.09 |
| 23 | P.En.U | 0.36 | -4.94 | -4.12 | -9.06 | -2.28 | 5.12 | 1.14 | 0.78 | 1.64 | 1.00 | 0.87 | 2.33 |
| 24 | P.En.F | 0.34 | -2.08 | -7.86 | -9.94 | -1.67 | 3.77 | 4.01 | 0.24 | 0.30 | 3.01 | 0.75 | 0.32 |
| 25 | A.En.U | 0.33 | -1.08 | -8.39 | -9.46 | -1.74 | 3.96 | 4.08 | 0.23 | 0.00 | 3.05 | 0.75 | 0.28 |
| 26 | P.En.U | 0.32 | -2.46 | -6.77 | -9.24 | -1.51 | 3.76 | 3.35 | 0.36 | 0.38 | 2.31 | 0.69 | 1.02 |
| 27 | P.En.F | 0.30 | -2.58 | -7.25 | -9.82 | -1.43 | 3.36 | 3.82 | 0.27 | 0.77 | 2.49 | 0.65 | 0.84 |
| 28 | P.En.F | 0.30 | -4.27 | -5.79 | -10.06 | -2.21 | 5.09 | 2.02 | 0.62 | 1.26 | 1.84 | 0.91 | 1.49 |
| 29 | A.En.U | 0.28 | -1.09 | -8.22 | -9.31 | -1.63 | 3.41 | 4.09 | 0.22 | 0.02 | 3.13 | 0.76 | 0.20 |

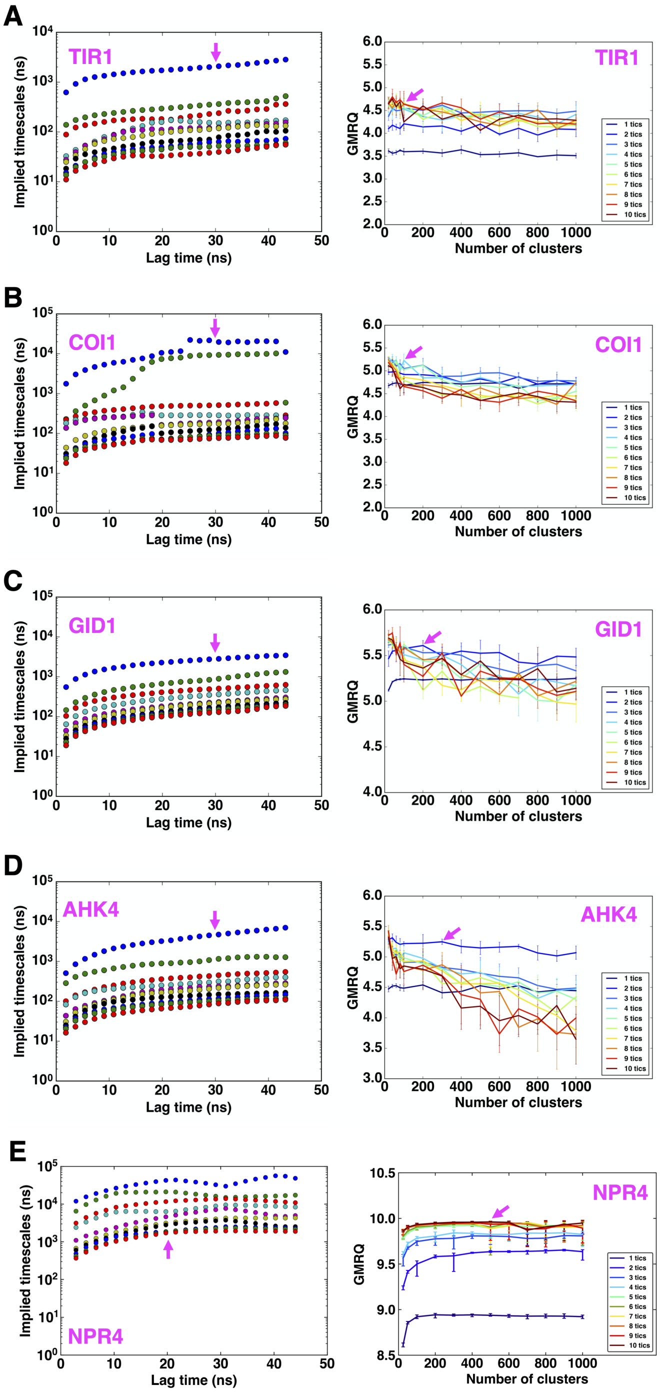


**Fig. S1.** Cross validation for Markov state models (MSMs) hyperparameter selection. Left panels: the slowest 10 implied timescales of MSMs constructed with increasing lag times converge after 20–30 ns for (A) IAA binding to TIR1, (B) JA-Ile binding to COI1, (C) GA3 binding to GID1, (D) trans-zeatin binding to AHK4, and (E) SA binding to NPR4. Right panels: the GMRQ scores for MSMs constructed with different number of clusters and number of tICs for each system (A–E). The scores were used to select the final MSMs used to describe the binding processes.

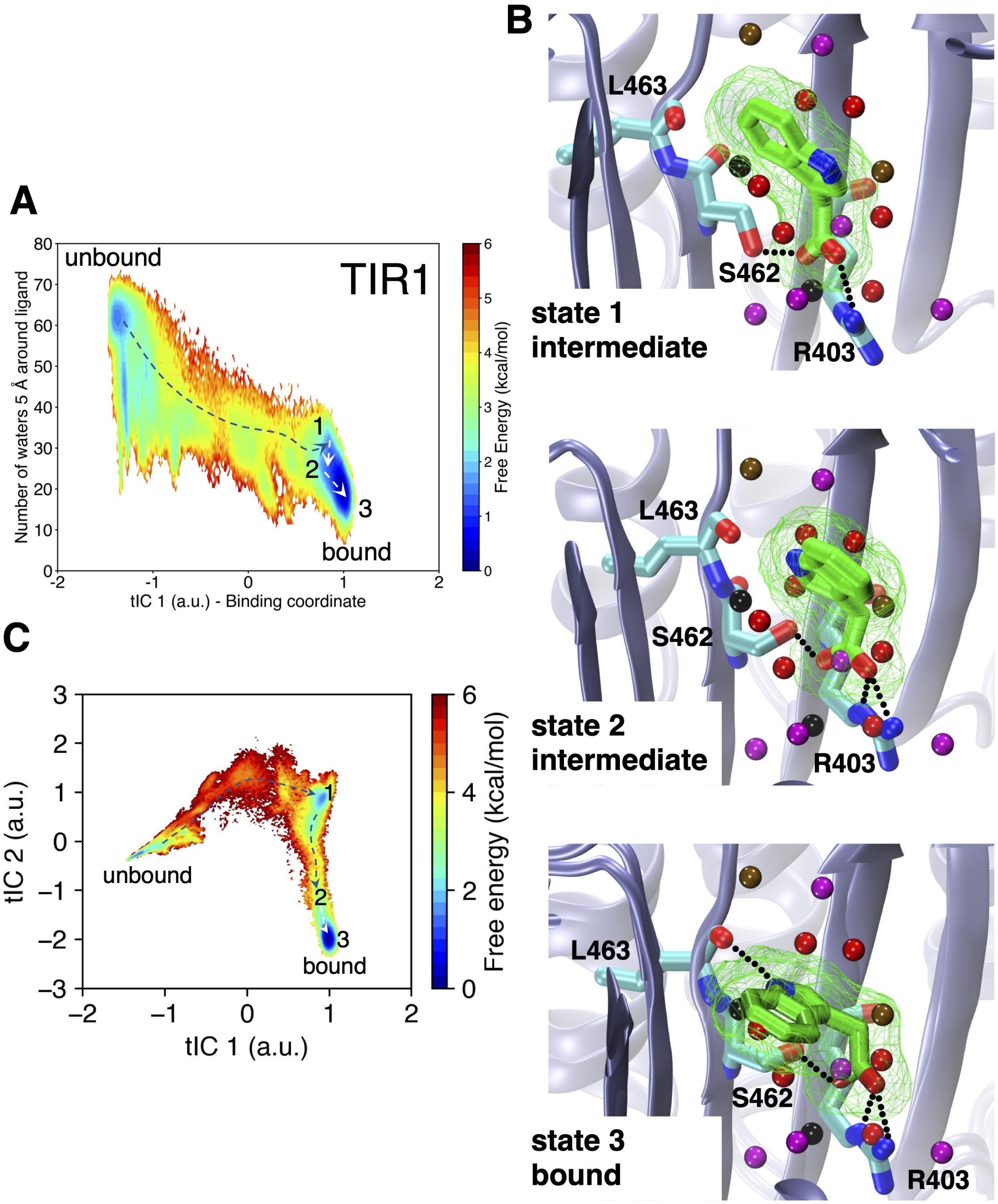


**Fig. S2.** (A) Free energy landscape of IAA binding to TIR1. The horizontal coordinate is the slowest process recovered from time-lagged independent component analysis (tICA) and is associated with ligand binding, while the vertical coordinate represents the ligand solvation. (B) The snapshots of the binding pathway for the binding of IAA to TIR1. The *apo* hydration sites are shown to indicate the exclusion of water molecules along the IAA binding process. (C) Free energy landscape of IAA binding to TIR1 in terms of the first two tICs.

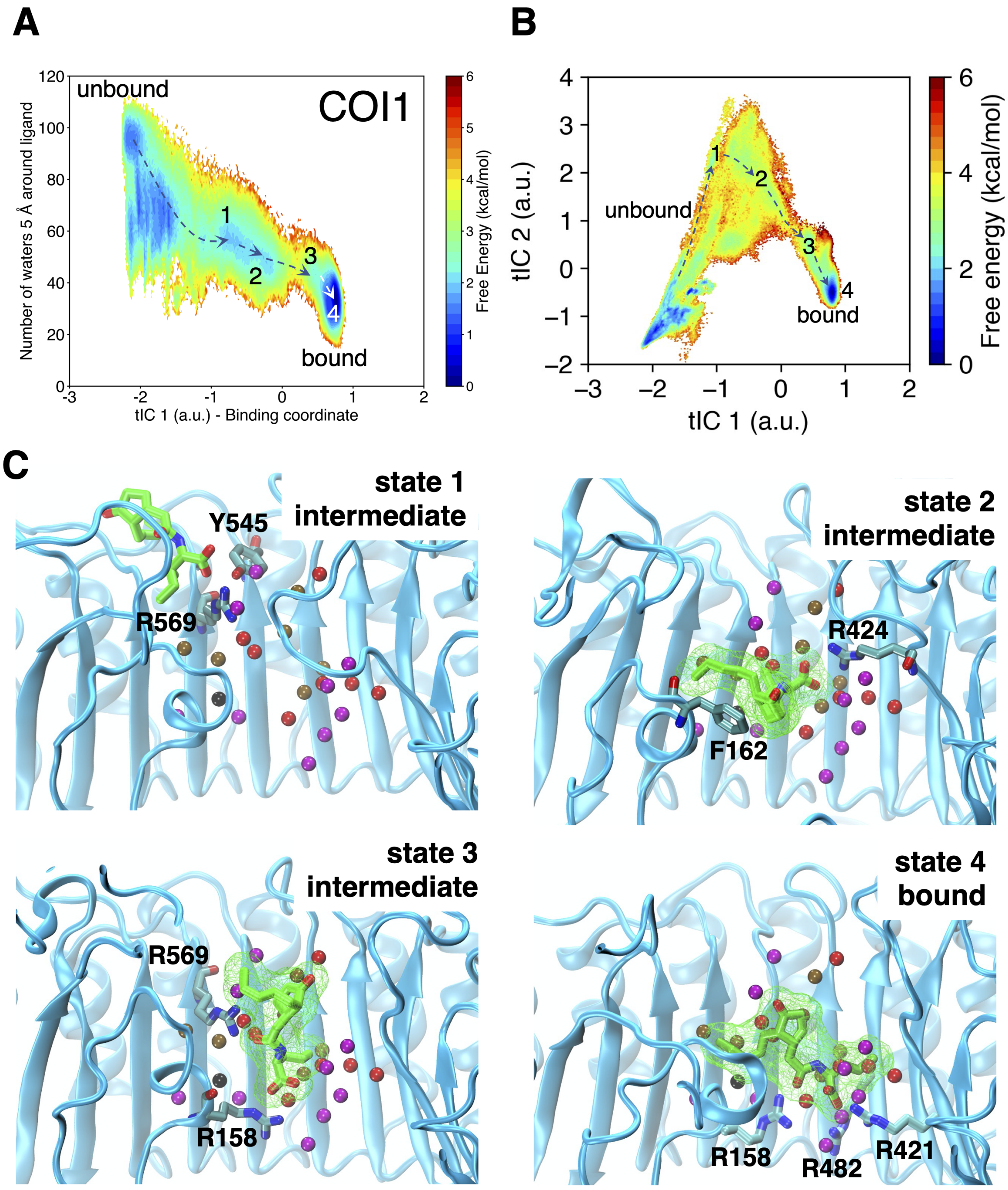


**Fig. S3.** (A) Free energy landscape of JA-Ile binding to COI1. The horizontal coordinate is the slowest process recovered from time-lagged independent component analysis (tICA) and is associated with ligand binding, while the vertical coordinate represents the ligand solvation. (B) Free energy landscape of JA-Ile binding to COI1 in terms of the first two tICs. (C) The snapshots of the binding pathway for the binding of JA-Ile to COI1. The *apo* hydration sites are shown to indicate the exclusion of water molecules along the JA-Ile binding process.

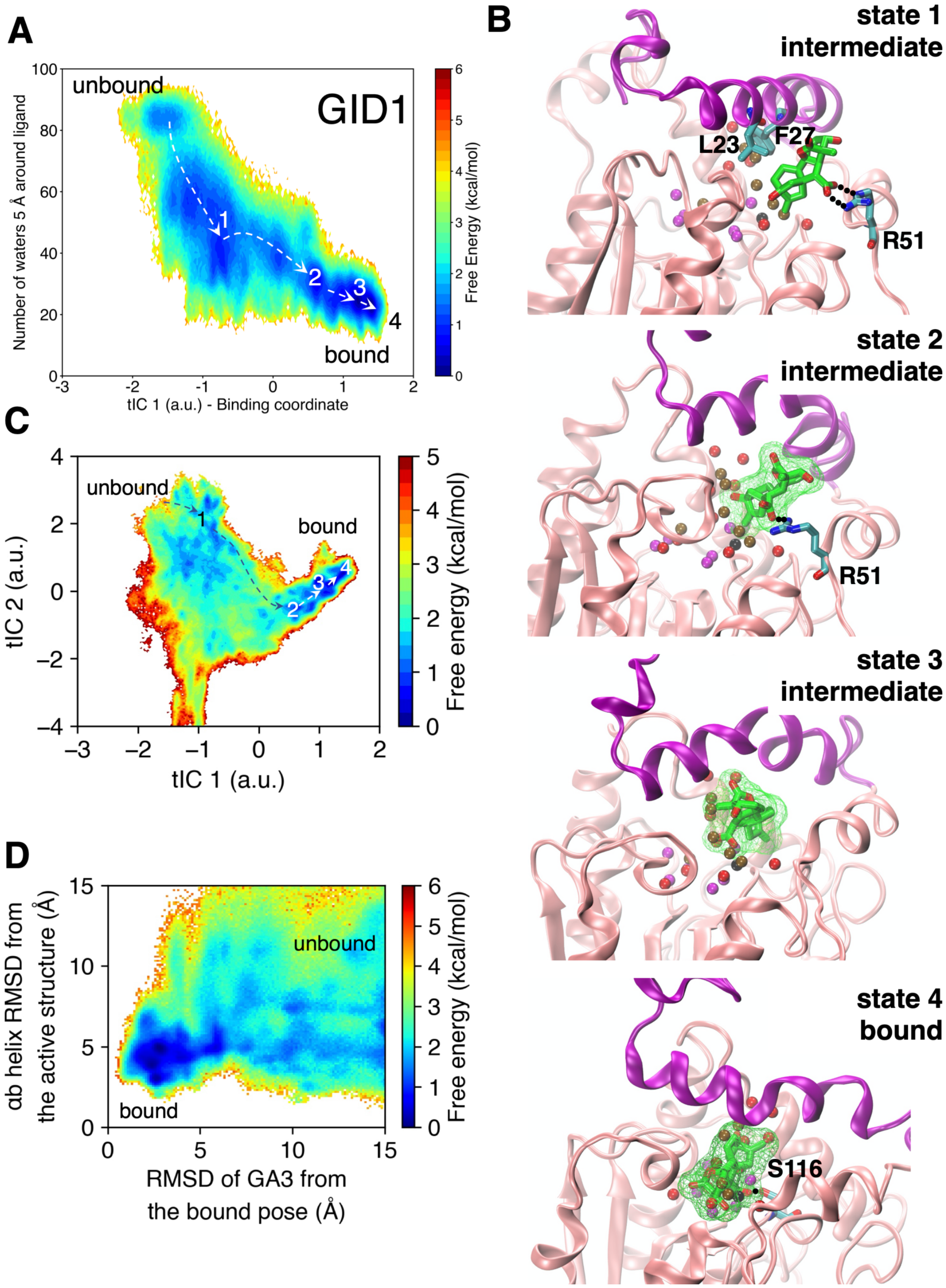


**Fig. S4.** (A) Free energy landscape of GA3 binding to GID1. The horizontal coordinate is the slowest process recovered from time-lagged independent component analysis (tICA) and is associated with ligand binding, while the vertical coordinate represents the ligand solvation. (B) The snapshots of the binding pathway for the binding of GA3 to GID1. The *apo* hydration sites are shown to indicate the exclusion of water molecules along the GA3 binding process. (C) Free energy landscape of GA3 binding to GID1 in terms of the first two tICs. (D) The binding of GA3 stabilizes the closure of $\alpha$b helix in the N-terminal of GID1.

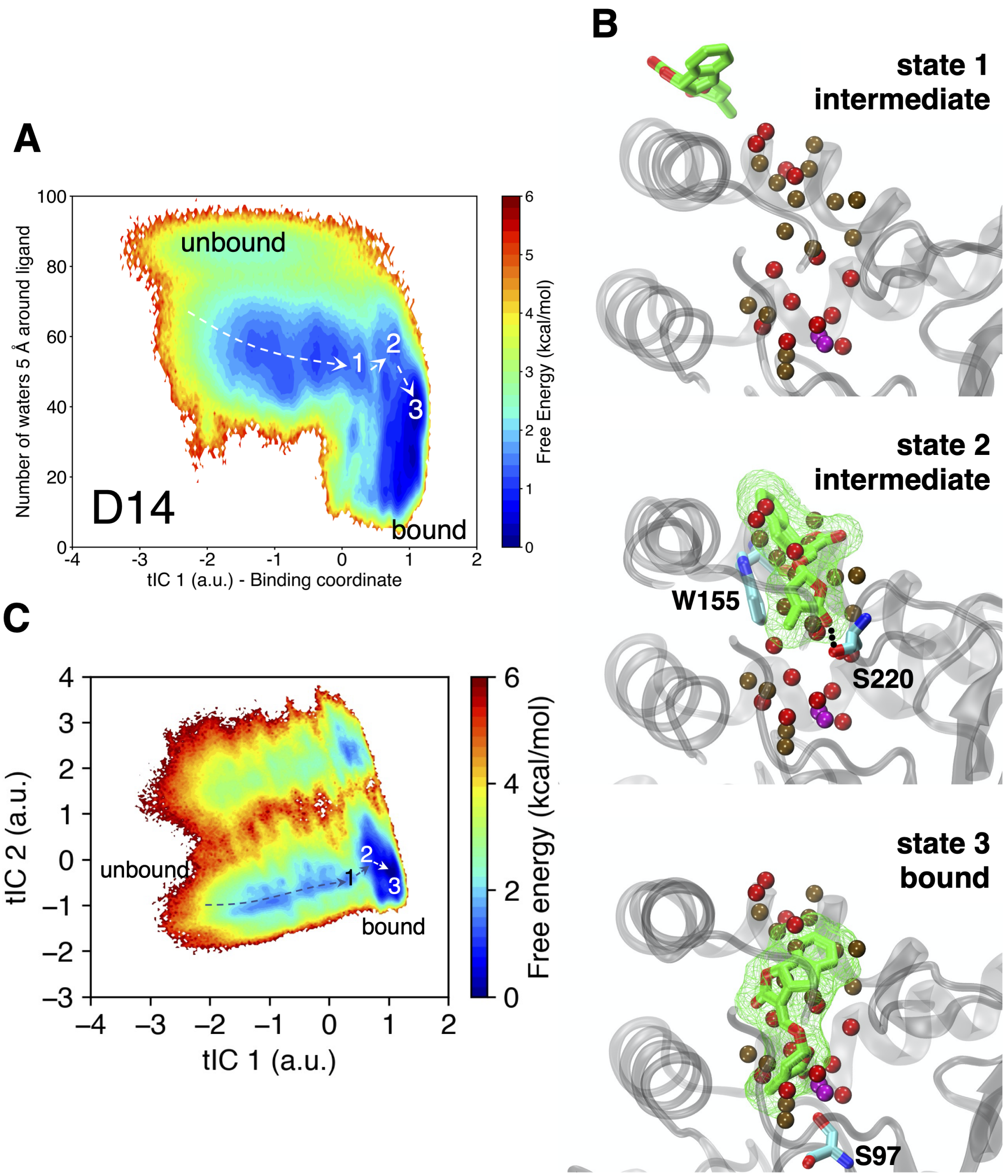


**Fig. S5.** (A) Free energy landscape of GR24 binding to D14. The horizontal coordinate is the slowest process recovered from time-lagged independent component analysis (tICA) and is associated with ligand binding, while the vertical coordinate represents the ligand solvation. (B) The snapshots of the binding pathway for the binding of GR24 to D14. The *apo* hydration sites are shown to indicate the exclusion of water molecules along the GR24 binding process. (C) Free energy landscape of GR24 binding to D14 in terms of the first two tICs.

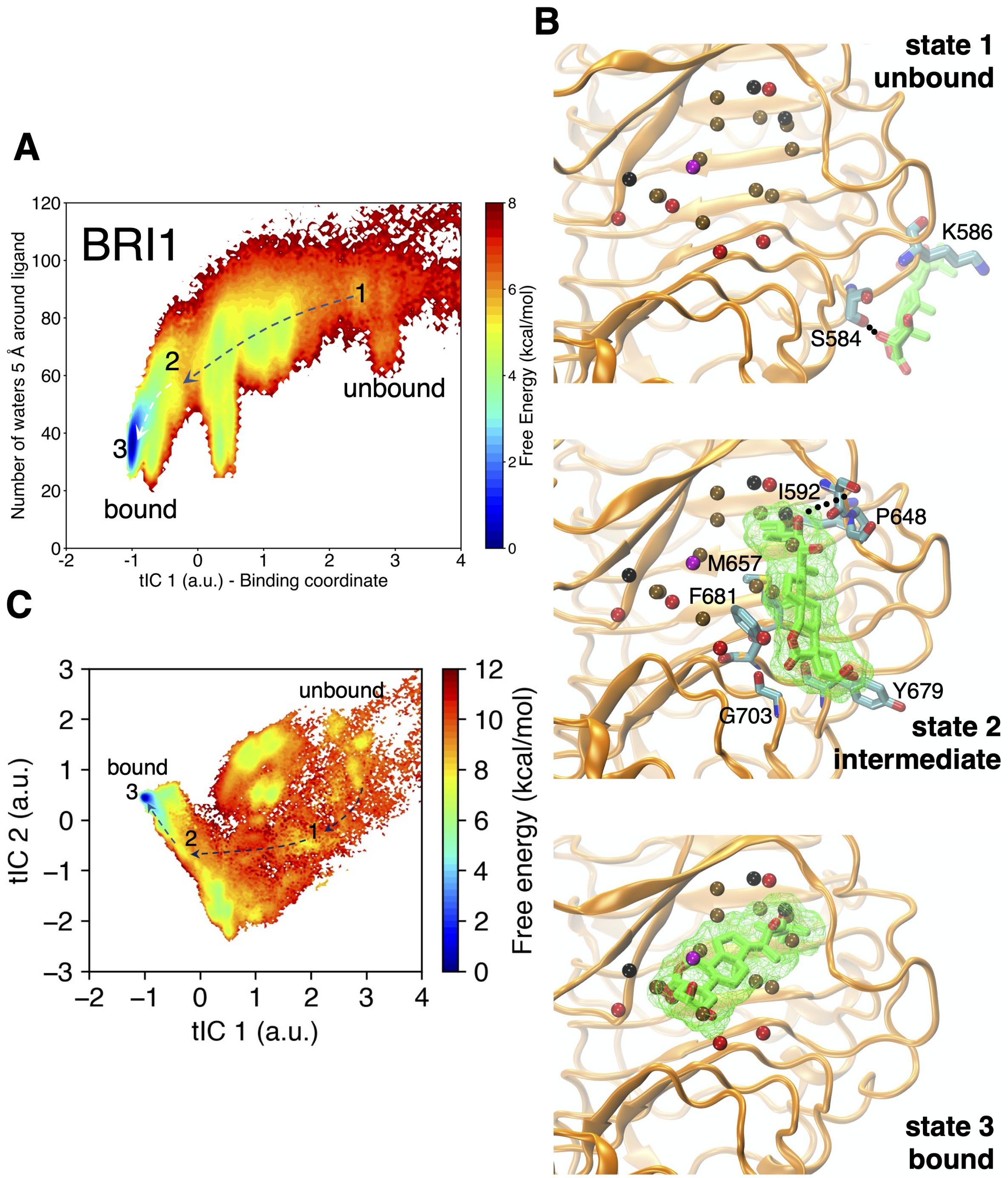


**Fig. S6.** (A) Free energy landscape of BLD binding to BRI1. The horizontal coordinate is the slowest process recovered from time-lagged independent component analysis (tICA) and is associated with ligand binding, while the vertical coordinate represents the ligand solvation. (B) The snapshots of the binding pathway for the binding of BLD to BRI1. The *apo* hydration sites are shown to indicate the exclusion of water molecules along the BLD binding process. (C) Free energy landscape of GR24 binding to D14 in terms of the first two tICs.

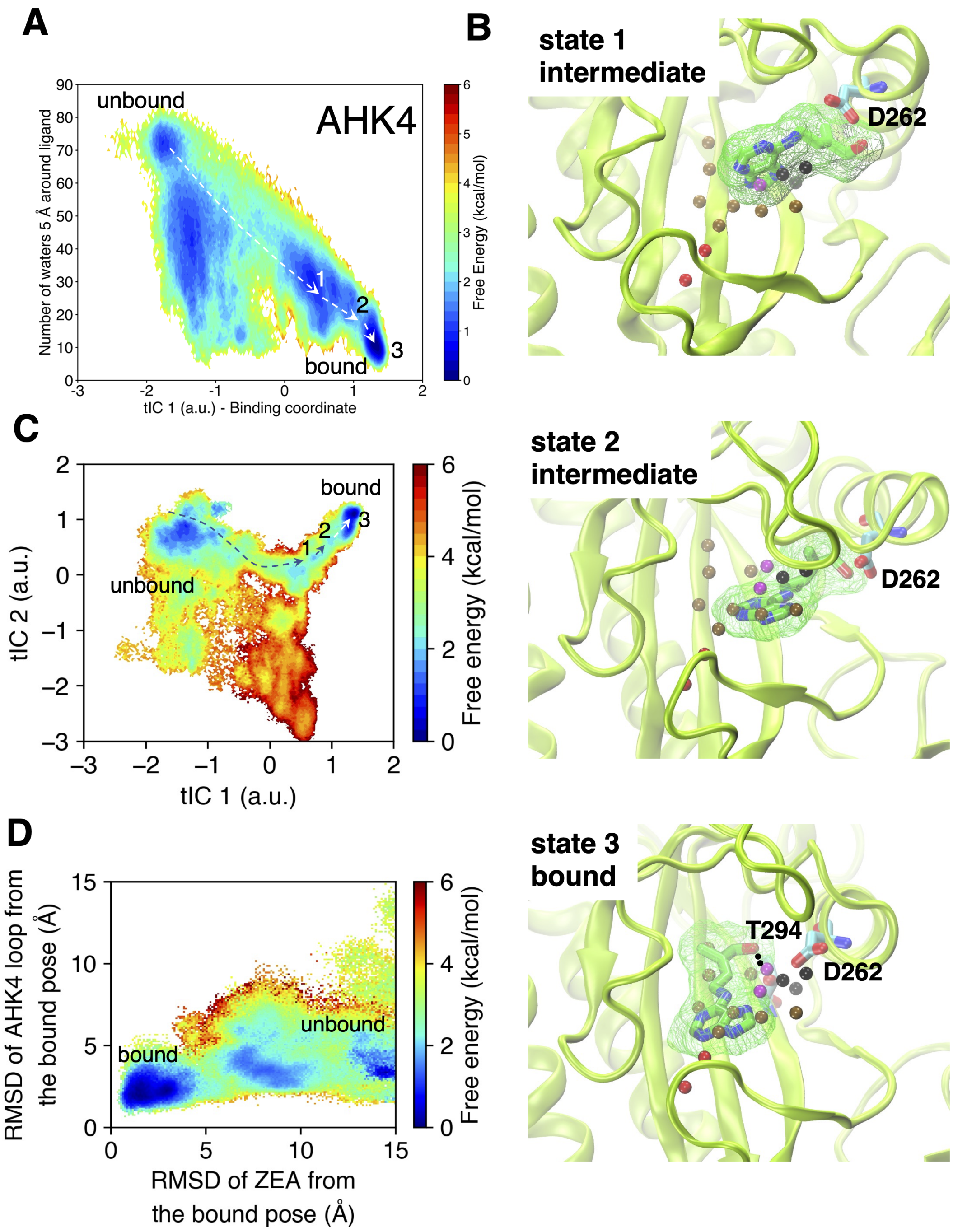


**Fig. S7.** (A) Free energy landscape of trans-Zeatin binding to AHK4. The horizontal coordinate is the slowest process recovered from time-lagged independent component analysis (tICA) and is associated with ligand binding, while the vertical coordinate represents the ligand solvation. (B) The snapshots of the binding pathway for the binding of trans-Zeatin to AHK4. The *apo* hydration sites are shown to indicate the exclusion of water molecules along the trans-Zeatin binding process. (C) Free energy landscape of trans-Zeatin binding to AHK4 in terms of the first two tICs. (D) The loop in AHK4 that encloses the binding site undergoes conformational changes during the binding of trans-Zeatin. The bound ligand stabilizes the loop after trans-Zeatin is bound to AHK4.

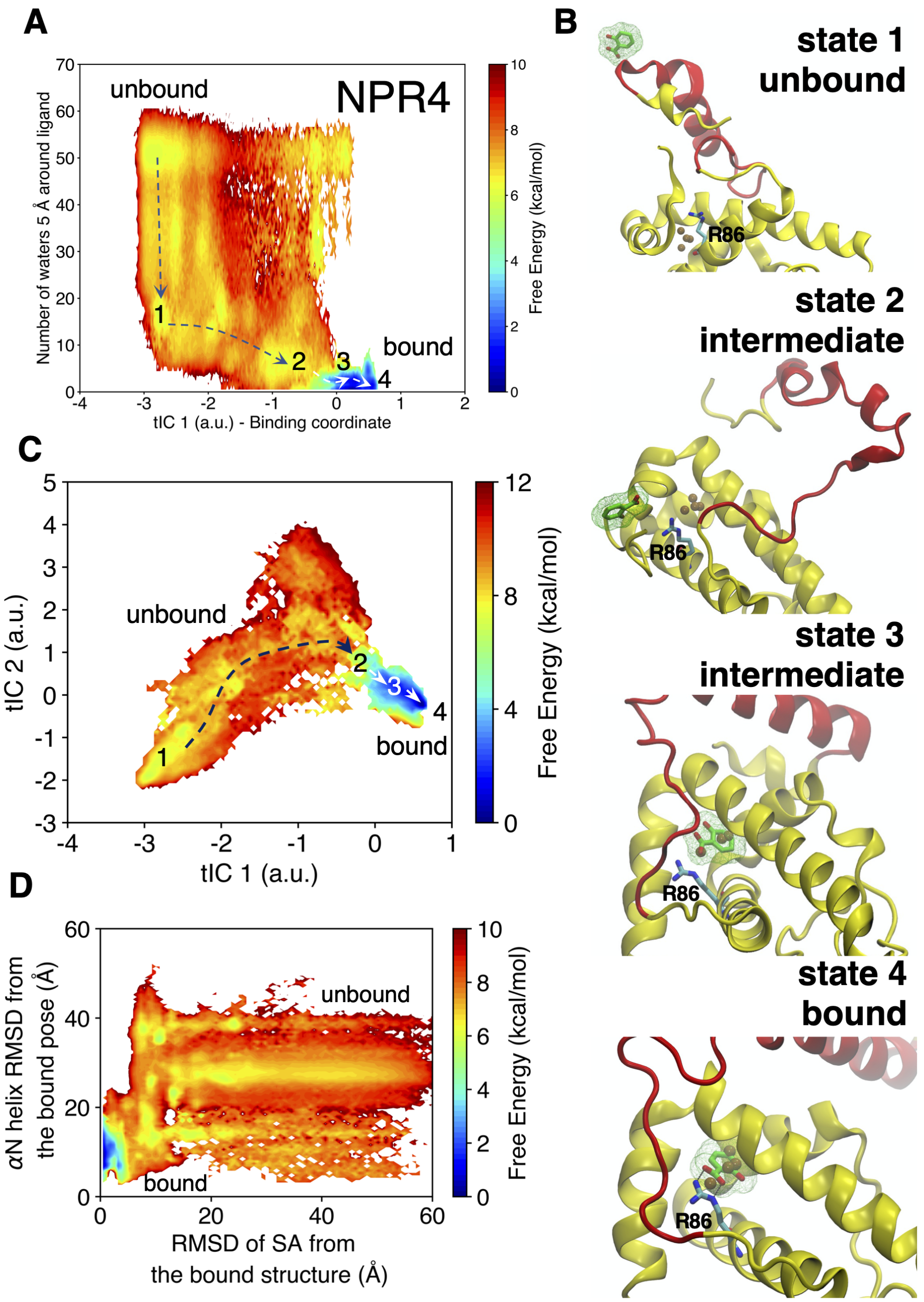


**Fig. S8.** (A) Free energy landscape of SA binding to the salicylic acid binding core (SBC) in NPR4. The horizontal coordinate is the slowest process recovered from time-lagged independent component analysis (tICA) and is associated with ligand binding, while the vertical coordinate represents the ligand solvation. (B) The snapshots of the binding pathway for the binding of SA to NPR4. The *apo* hydration sites are shown to indicate the exclusion of water molecules along the SA binding process. Unlike other phytohormones, SA displaces all the water molecules from the pocket upon ligand binding. (D) The binding of SA likely stabilizes the closure of the $\alpha$N helix and loop (shown in red) in the N-terminal of the SBC.

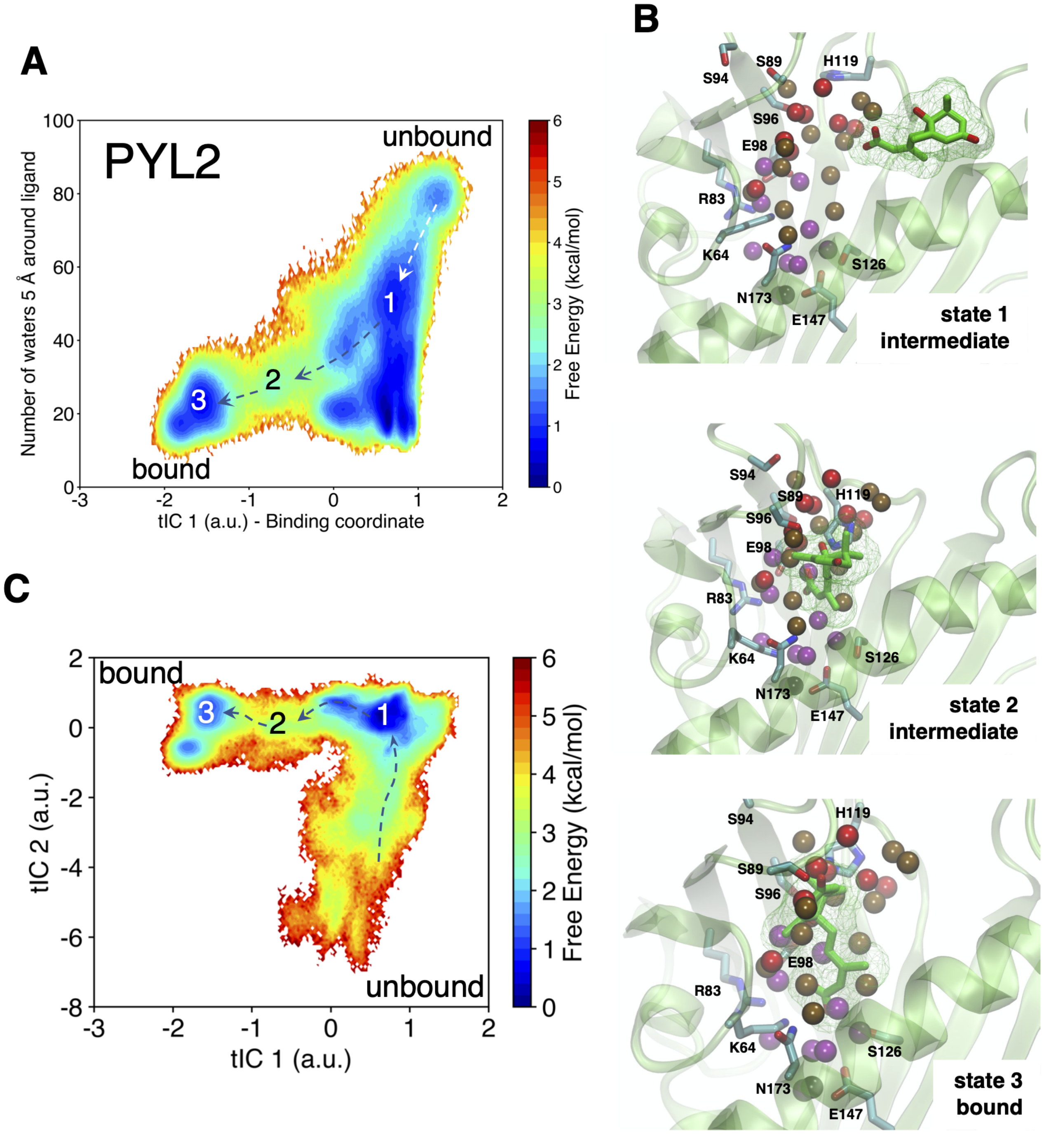


**Fig. S9.** (A) Free energy landscape of ABA binding to PYL2. The horizontal coordinate is the slowest process recovered from time-lagged independent component analysis (tICA) and is associated with ligand binding, while the vertical coordinate represents the ligand solvation. (B) The snapshots of the binding pathway for the binding of abscisic acid (ABA) to PYL2. The *apo* hydration sites are shown to indicate the exclusion of water molecules along the ABA binding process. (C) Free energy landscape of ABA binding to PYL2 in terms of the first two tICs.

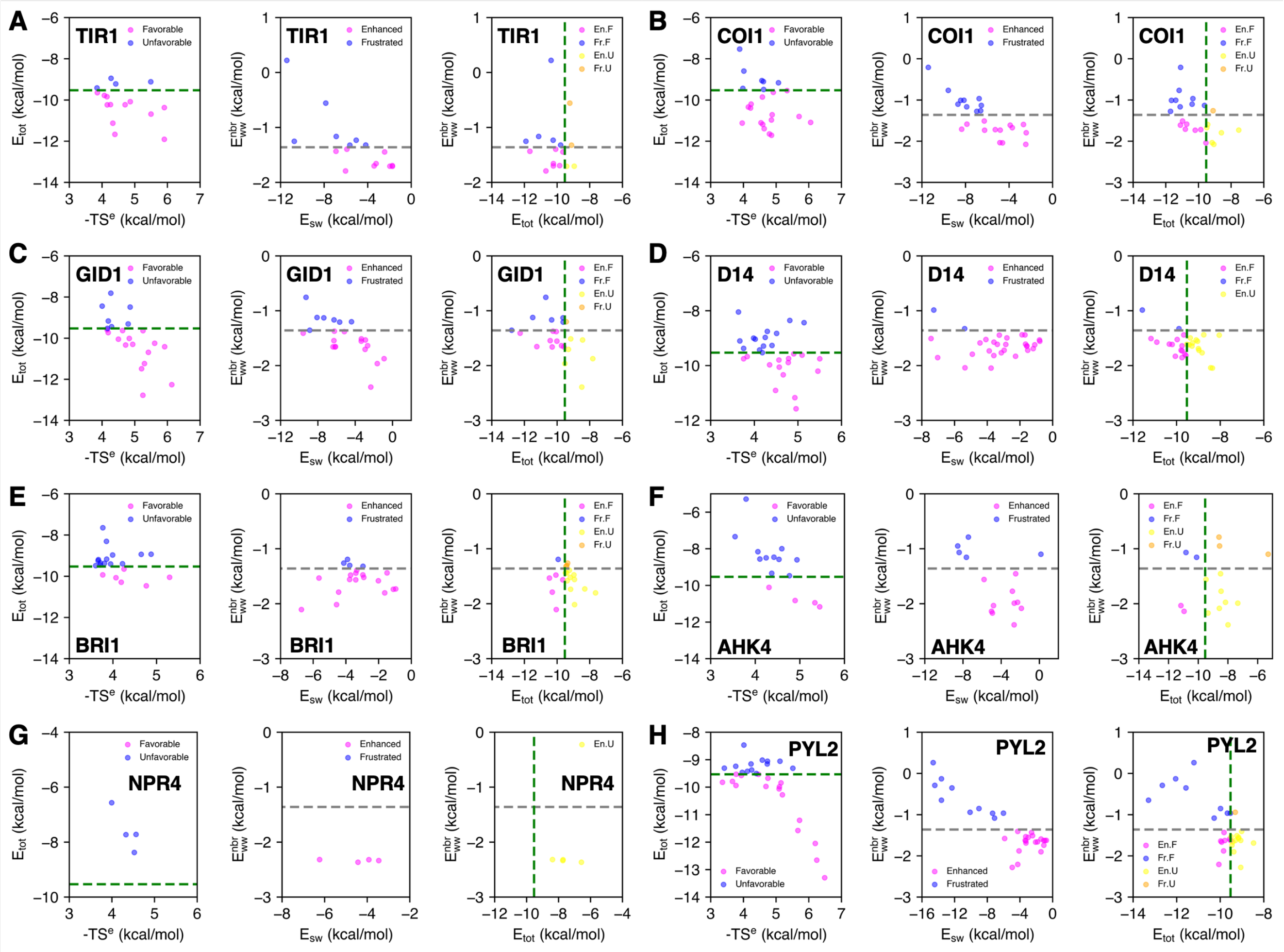

**Fig. S10.** Thermodynamic properties of *apo* hydration sites in (A) TIR1, (B) COI1, (C) GID1, (D) D14, (E) BRI1, (F) AHK4, (G) NPR4, and (H) PYL2. (Left) $E_{tot}$ and -TS$^e$ of the favorable (magenta) and unfavorable (blue) hydration sites, (middle) $E_{ww}^{nbr}$ and $E_{sw}$ of the enhanced (magenta) and frustrated (blue) hydration sites, and (right) $E_{ww}^{nbr}$ and $E_{tot}$ of En.F (magenta), Fr.F (blue), En.U (yellow) and Fr.U (orange) hydration sites.

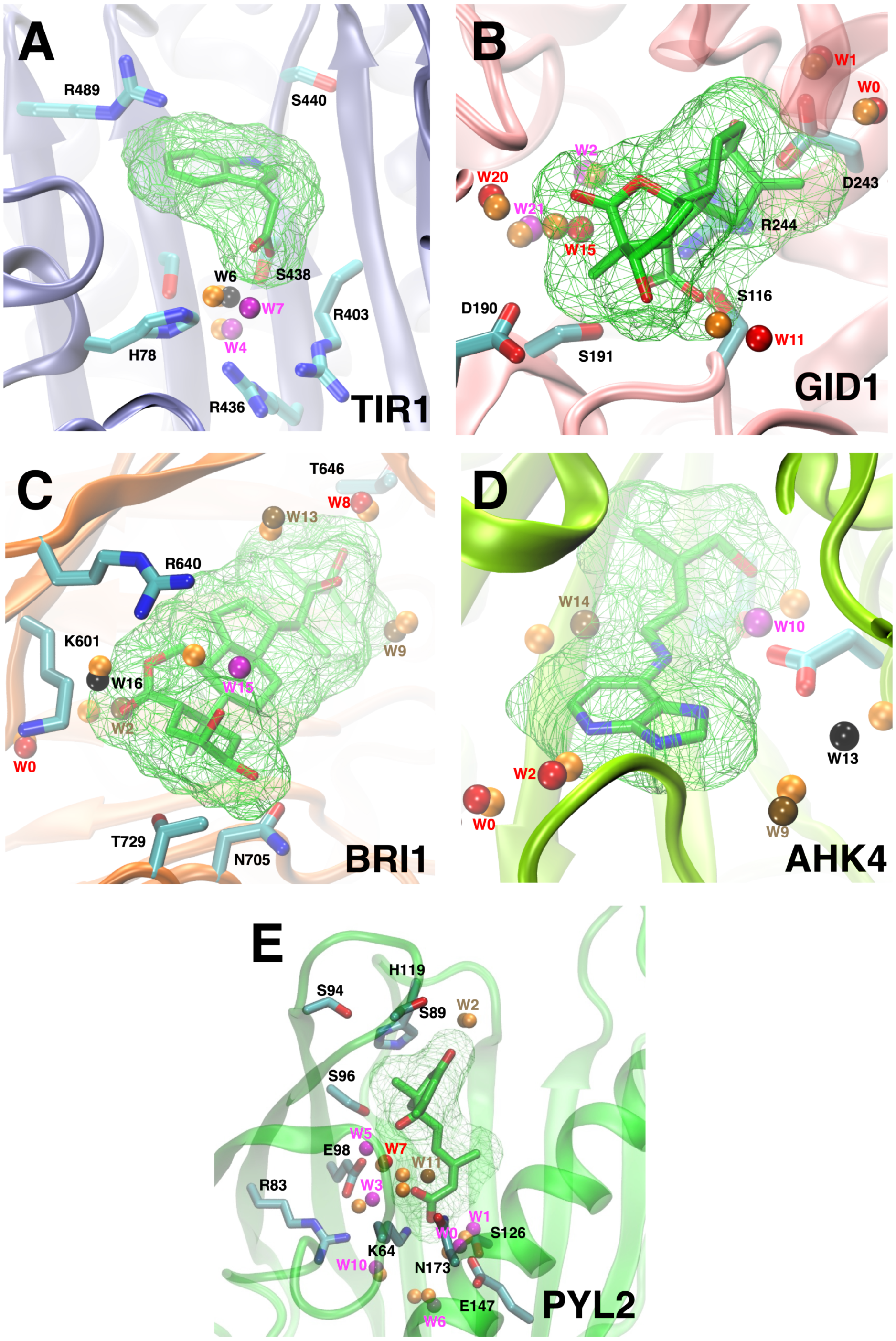


**Fig. S11.** Overlay of *apo* hydration sites identified from MD simulations (En.F: red, En.U: ochre, Fr.F: magenta, Fr.U: black) and water molecules captured in the crystal structure (orange) of the receptor-hormone bound complexes. (A) TIR1, (B) GID1, (C) BRI1, (D) AHK4, and (E) PYL2.

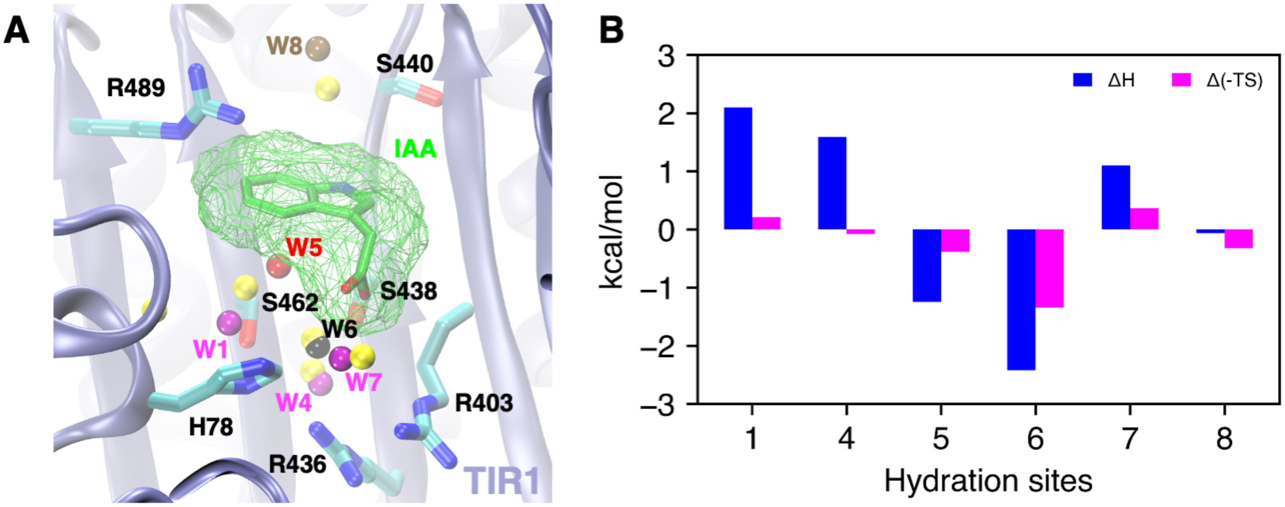


**Fig. S12.** (A) Overlay of the conserved *apo* hydration sites (labelled) and the corresponding *holo* sites (yellow) in the binding site of TIR1 upon the binding of IAA, and (B) the changes in enthalpy and entropy of conserved hydration sites.

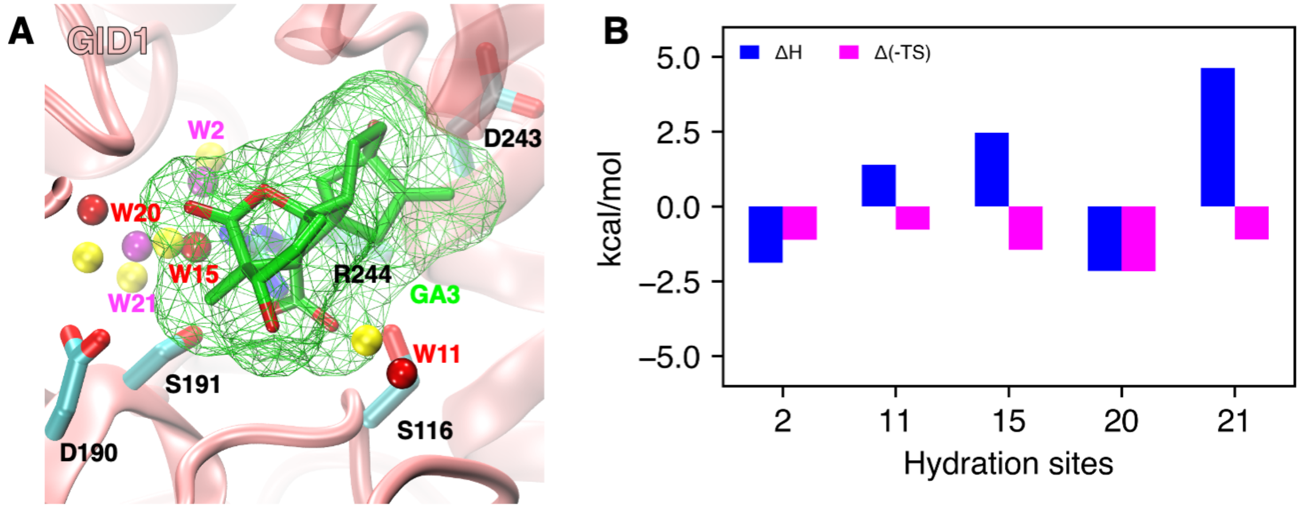


**Fig. S13.** (A) Overlay of the conserved *apo* hydration sites (labelled) and the corresponding *holo* sites (yellow) in the binding site of GID1 upon the binding of GA3, and (B) the changes in enthalpy and entropy of conserved hydration sites.

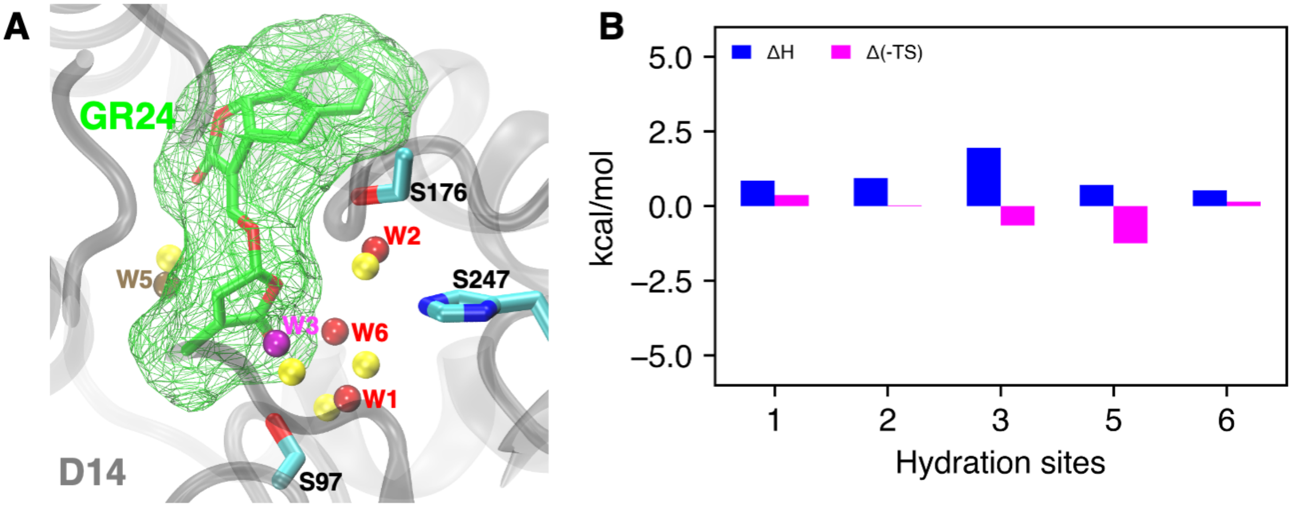


**Fig. S14.** (A) Overlay of the conserved *apo* hydration sites (labelled) and the corresponding *holo* sites (yellow) in the binding site of D14 upon the binding of GR24, and (B) the changes in enthalpy and entropy of conserved hydration sites.

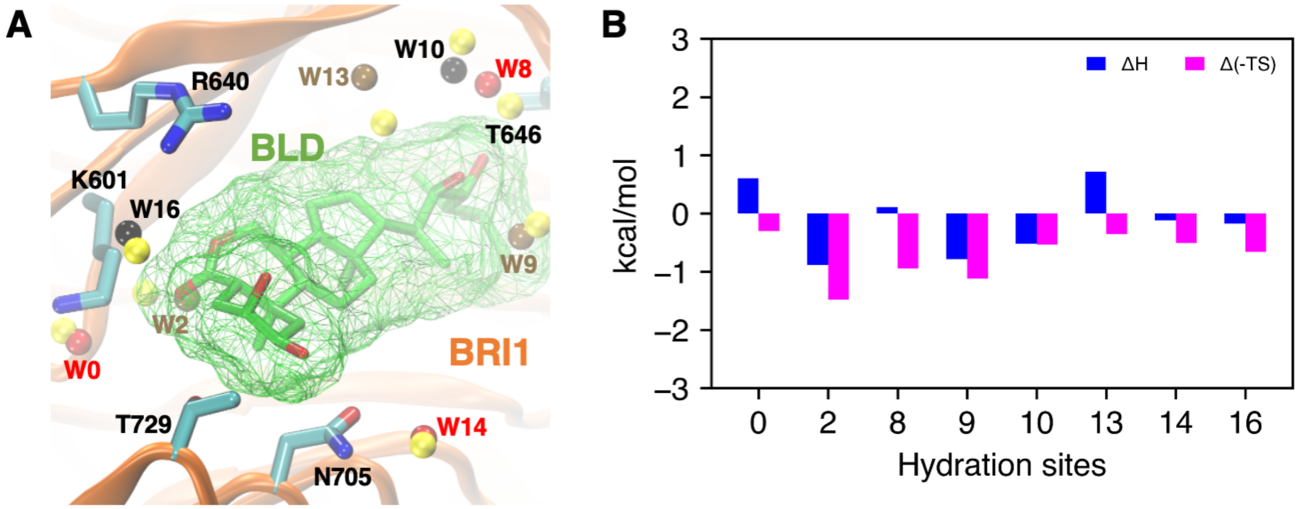


**Fig. S15.** (A) Overlay of the conserved *apo* hydration sites (labelled) and the corresponding *holo* sites (yellow) in the binding site of BRI1 upon the binding of GR24, and (B) the changes in enthalpy and entropy of conserved hydration sites.

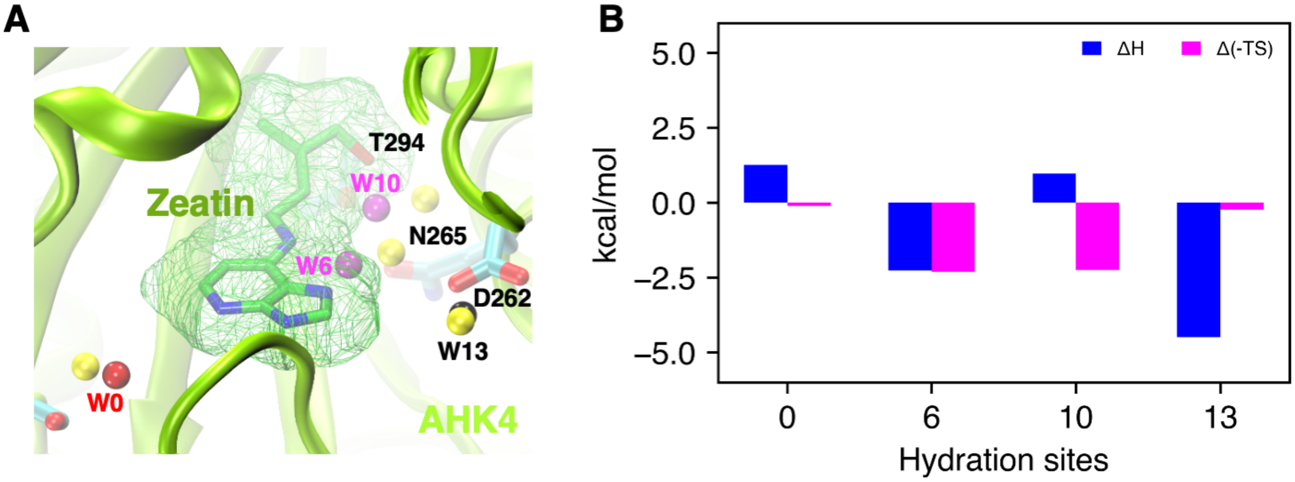


**Fig. S16.** (A) Overlay of the conserved *apo* hydration sites (labelled) and the corresponding *holo* sites (yellow) in the binding site of AHK4 upon the binding of trans-Zeatin, and (B) the changes in enthalpy and entropy of conserved hydration sites.

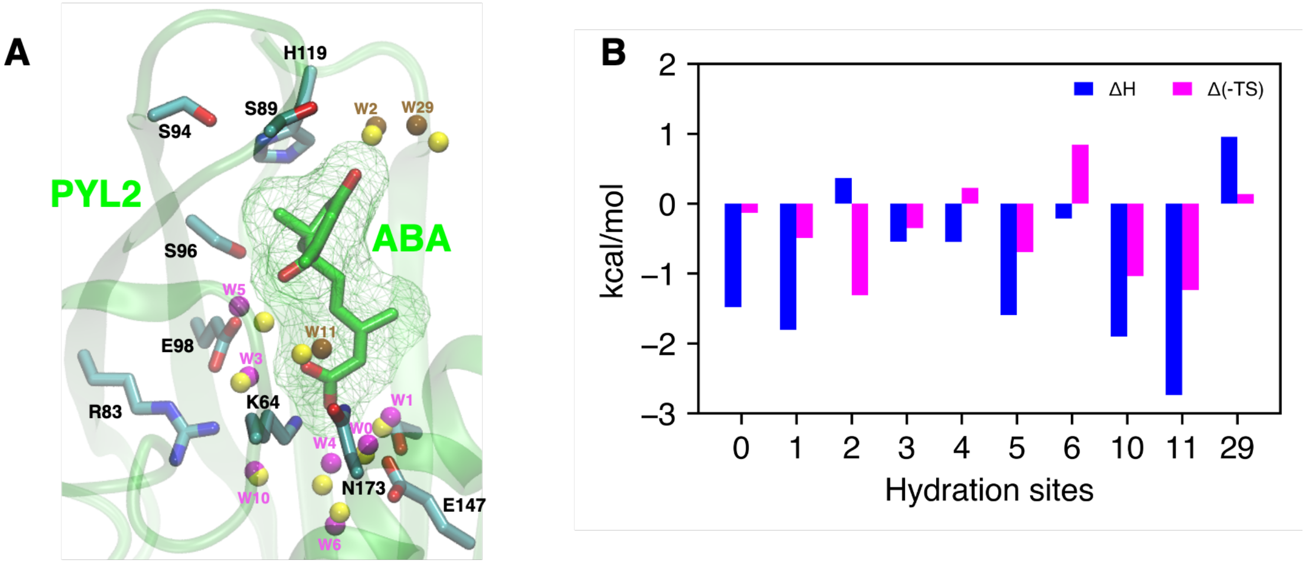


**Fig. S17.** (A) Overlay of the conserved *apo* hydration sites (labelled) and the corresponding *holo* sites (yellow) in the binding site of PYL2 upon the binding of ABA, and (B) the changes in enthalpy and entropy of conserved hydration sites.

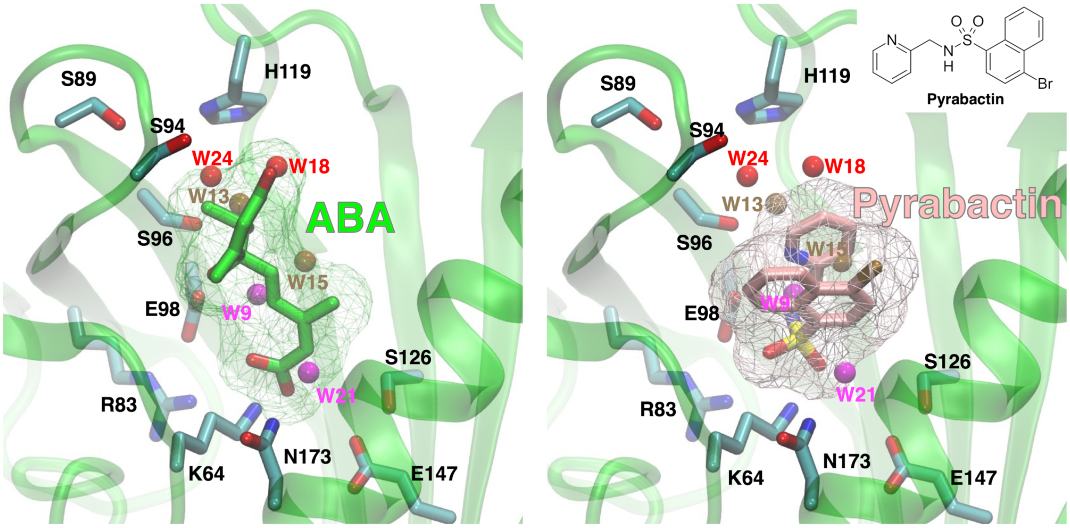


**Fig. S18.** Hydration site analysis reveals different waters are displaced by ABA and pyrabactin upon binding to PYL2. Left: overlay of ABA and *apo* hydration sites in PYL2. Right: overlay of pyrabactin and *apo* hydration sites in PYL2.

### References


[1] A. Waterhouse, M. Bertoni, S. Bienert, G. Studer, G. Tauriello, R. Gumienny, F. T. Heer, T. A. P. de Beer, C. Rempfer, L. Bordoli, R. Lepore and T. Schwede, *Nucleic Acids Research*, 2018, **46**, W296–W303.

[2] A. Sali, *Molecular Medicine Today*, 1995, **1**, 270–277.

[3] M. yi Shen and A. Sali, *Protein Science*, 2006, **15**, 2507–2524.

[4] L. Martínez, R. Andrade, E. G. Birgin and J. M. Martínez, *J. Comput. Chem.*, 2009, **30**, 2157–2164.

[5] J. Wang, R. M. Wolf, J. W. Caldwell, P. A. Kollman and D. A. Case, *J. Comput. Chem.*, 2004, **25**, 1157–1174.

[6] S. A. J. Rosen, P. R. J. Gaffney and I. R. Gould, *Phys. Chem. Chem. Phys.*, 2011, **13**, 1070–1081.

[7] T. Darden, D. York and L. Pedersen, *J. Chem. Phys.*, 1993, **98**, 10089–10092.

[8] J.-P. Ryckaert, G. Ciccotti and H. J. Berendsen, *J. Comput. Phys.*, 1977, **23**, 327–341.

[9] L. C. Pierce, R. Salomon-Ferrer, C. A. F. de Oliveira, J. A. McCammon and R. C. Walker, *Journal of Chemical Theory and Computation*, 2012, **8**, 2997–3002.

[10] Y. Naritomi and S. Fuchigami, *J. Chem. Phys.*, 2011, **134**, 065101.

[11] R. T. McGibbon and V. S. Pande, *J. Chem. Phys.*, 2015, **142**, 124105.

[12] M. P. Harrigan, M. M. Sultan, C. X. Hernández, B. E. Husic, P. Eastman, C. R. Schwantes, K. A. Beauchamp, R. T. McGibbon and V. S. Pande, *Biophys. J.*, 2017, **112**, 10–15.

[13] R. T. McGibbon, C. X. Hernández, M. P. Harrigan, S. Kearnes, M. M. Sultan, S. Jastrzebski, B. E. Husic and V. S. Pande, *J. Open Source Software*, 2016, **1**, 00034.

[14] F. Aldukhi, A. Deb, C. Zhao, A. S. Moffett and D. Shukla, *J. Phys. Chem. B*, 2019, **124**, 355–365.

[15] J. Chen, A. White, D. C. Nelson and D. Shukla, *J. Biol. Chem.*, 2021, **297**, 101092.

[16] C. Zhao and D. Shukla, *bioRxiv, doi: 10.1101/721761*, 2021.

[17] W. E and E. Vanden-Eijnden, *J. Stat. Phys.*, 2006, **123**, 503.

[18] W. E and E. Vanden-Eijnden, *Annu. Rev. Phys. Chem.*, 2010, **61**, 391–420.

[19] I. Y. Ben-Shalom, C. Lin, T. Kurtzman, R. C. Walker and M. K. Gilson, *J. Chem. Theory Comput.*, 2019, **15**, 2684–2691.

[20] K. Haider, A. Cruz, S. Ramsey, M. K. Gilson and T. Kurtzman, *J. Chem. Theory Comput.*, 2017, **14**, 418–425.

[21] R. Abel, T. Young, R. Farid, B. J. Berne and R. A. Friesner, *J. Am. Chem. Soc.*, 2008, **130**, 2817–2831.

[22] T. Lazaridis, *J. Phys. Chem. B*, 1998, **102**, 3531–3541.
